# Microstates and power envelope hidden Markov modeling probe bursting brain activity at different timescales

**DOI:** 10.1101/2021.02.20.432128

**Authors:** N Coquelet, X De Tiège, L Roshchupkina, P Peigneux, S Goldman, M Woolrich, V Wens

## Abstract

State modeling of whole-brain electroencephalography (EEG) or magnetoencephalography (MEG) allows to investigate transient, recurring neurodynamical events. Two widely-used techniques are the microstate analysis of EEG signals and hidden Markov modeling (HMM) of MEG power envelopes. Both reportedly lead to similar state lifetimes on the 100 ms timescale, suggesting a common neural basis. We addressed this issue by using simultaneous MEG/EEG recordings at rest and comparing the spatial signature and temporal activation dynamics of microstates and power envelope HMM states obtained separately from EEG and MEG. Results showed that microstates and power envelope HMM states differed both spatially and temporally. Microstates tend to exhibit spatio-temporal locality, whereas power envelope HMM states disclose network-level activity with 100–200 ms lifetimes. Further, MEG microstates do not correspond to the canonical EEG microstates but are better interpreted as split HMM states. On the other hand, both MEG and EEG HMM states involve the (de)activation of similar functional networks. Microstate analysis and power envelope HMM thus appear sensitive to neural events occurring over different spatial and temporal scales. As such, they represent complementary approaches to explore the fast, sub-second scale bursting electrophysiological dynamics in spontaneous human brain activity.

## 1. Introduction

A fundamental part of human neural dynamics is the spontaneous emergence of brain rhythms, i.e., large-scale oscillations of neuroelectric activity (for a review, see, e.g., (Hari and Salmelin, 1997)). These rhythms play a critical role for human brain functions such as sensory, motor and cognitive processes ((Klimesch, 2012), for reviews, see (Klimesch et al., 2010; Pfurtscheller and Lopes da Silva, 1999)). They also wax and wane spontaneously at rest (i.e., in the absence of any explicit task performance). The resulting fluctuations in their amplitude are key to intrinsic functional connectivity (Siegel et al., 2012; Sjøgård et al., 2020a). When measured with electroencephalography (EEG) or magnetoencephalography (MEG), this oscillatory dynamics leads to signal power time courses whose correlation structure identifies functional brain networks (Brookes et al., 2011; Coquelet et al., 2020a; Hipp et al., 2012; Liu et al., 2017; Siems et al., 2016; Wens et al., 2014). Further, spontaneous MEG/EEG power fluctuations occur in transient, sub-second long bursts of oscillatory activity (van Ede et al., 2018). Short-lived power bursts may actually correspond to the fast activation/deactivation of functional networks (Baker et al., 2014; Britz et al., 2010; Vidaurre et al., 2018) and their co-occurrence, to the intrinsic functional connectivity of these networks (Seedat et al., 2020). They might ultimately relate to the metastable cross-network interactions characteristic of functional integration at the supra-second timescale (de Pasquale et al., 2016, 2012; Della Penna et al., 2019; Wens et al., 2019). Power bursts also presumably hold specific functions, such as the encoding of recently acquired information by coactivation with spontaneous replays (Higgins et al., 2020). Exploring the spontaneous dynamics of MEG/EEG power bursts thus represents a fundamental step towards a better understanding of the intrinsic functional architecture of the human brain.

With their millisecond-scale temporal resolution, EEG and MEG (Hari and Puce, 2017) are natural techniques to investigate power bursts, although the role of short-time events has also been emphasized with functional magnetic resonance imaging (fMRI) (Tagliazucchi et al., 2012). Accordingly, the two main data-driven methods used to detect recurring events of high electrophysiological power are EEG microstate analysis ((Lehmann et al., 1987); for a review, see (Michel and Koenig, 2018)) and hidden Markov modeling (HMM) of MEG power envelopes (Baker et al., 2014; Quinn et al., 2018). Both allow to partition EEG/MEG data into discrete brain states that recurrently activate and deactivate one after the other, yet the underlying clustering algorithms strongly differ in their assumptions and methods. Microstates are determined as time periods of quasi-stable scalp EEG topography that repeatedly occur, up to amplitude rescalings and polarity flips. Four canonical microstates have been identified with reported mean lifetimes ranging from 60 to 120 ms (for a review, see, e.g. (Michel and Koenig, 2018)). These microstates were associated with different classes of mentation (Lehmann et al., 1998) and partially correlated with the spontaneous haemodynamics of some fMRI networks (Britz et al., 2010; Musso et al., 2010; Yuan et al., 2012). Their temporal properties are also affected by brain disorders such as schizophrenia (Koenig et al., 1999; Lehmann et al., 2005) or multiple sclerosis (Gschwind et al., 2016). By contrast, the HMM relies on the more abstract concept of Markov chains to describe brain power dynamics in terms of causal transitions among “hidden” states (Rabiner, 1989). These states are hidden in the sense that they are not explicitly expressed in the data and must be inferred through implicit statistical features such as, e.g., the covariance matrix of a state observation model (Rezek and Roberts, 2005). Here, the HMM states are determined by transient patterns of MEG power envelope covariance repeating over time (Baker et al., 2014), but occurring on too short time periods to be measurable directly from the data, e.g., with sliding windows (for a review, see (O’Neill et al., 2018)). The HMM inference applied to MEG power envelope signals has typically been used to identify 6 or 8 states disclosing a spatial distribution reminiscent of brain functional networks as well as mean lifetimes ranging from 50 to 200 ms (Baker et al., 2014; Quinn et al., 2018). Temporal properties of HMM states are also altered by physiological processes such as ageing (Brookes et al., 2018; Coquelet et al., 2020b) as well as brain disorders such as Alzheimer’s disease (Puttaert et al., 2020; Sitnikova et al., 2018) or multiple sclerosis (Van Schependom et al., 2019).

Interestingly, despite fundamental methodological differences, EEG microstates and MEG power envelope HMM states appear to remain stable over similar timescales. This raises the question of whether they describe similar neurodynamics ((Baker et al., 2014); for a review, see (Khanna et al., 2015)). Here, we investigate this key question using simultaneous MEG/EEG recordings of resting-state activity. Still, comparing EEG microstates and MEG power envelope HMM states entangles two potential issues, i.e., the effect of the state clustering model (microstates vs. HMM) and that of the recording modality (EEG vs. MEG). To avoid such confound, we sought to adapt the notion of microstates to MEG, and that of HMM states to EEG, before conducting the comparison. To the best of our knowledge, a microstate analysis of MEG data has not yet been developed. The HMM approach has been applied to EEG power envelopes (Hunyadi et al., 2019; Sitnikova et al., 2020), but the focus was on the relationship with fMRI networks rather than MEG or EEG microstates. Here, we assessed the impact of both the state clustering model and the recording modality on temporal and spatial signatures of transient brain states. More specifically, we estimated to what extent two types of states tend to co-activate by temporal correlation analysis of their activation dynamics, and to what extent they involve similar brain regions or networks by spatial correlation analysis of the associated power distributions. Based on the idea that microstates and HMM states are both designed to identify discrete recurrent brain states and given the reported similarity of their typical lifetimes (Baker et al., 2014), we hypothesized that the two state clustering models would reveal a close spatio-temporal relationship within each recording modality. This would suggest that microstates and power envelope HMM states disclose similar neural events. On the other hand, based on a previous comparison of MEG and EEG power envelope signals at rest (Coquelet et al., 2020a), we expected similar spatial signatures but substantially different temporal state dynamics across the two recording modalities.

## 2. Methods

### 2.1. Participants

Forty-two young adults (14 females, mean age ± standard deviation (SD): 24.4 ± 3.9 years, range: 18–35 years) were included in this study, 19 of which were already used in a previous study of our group (Coquelet et al., 2020a). All participants were right-handed according to the Edinburgh handedness inventory (Oldfield, 1971), did not take any psychotropic drug, and had no prior history of neurological or psychiatric disorder. Each of them signed a written informed consent before scanning. The CUB – Hôpital Erasme Ethics Committee approved this study prior to their inclusion.

### 2.2. Data acquisition

Participants underwent a resting-state recording session (eyes open, fixation cross, 5 minutes) with simultaneous MEG and high-density EEG. Neuromagnetic activity was recorded with a 306-channel whole-scalp MEG system (band-pass: 0.1–330 Hz, sampling frequency: 1 kHz) installed in a light-weight magnetically shielded room (Maxshield™, MEGIN, Cronton Healthcare, Helsinki, Finland; see (De Tiège et al., 2008) for detailed characteristics). Four coils continuously tracked subjects’ head position inside the MEG helmet. The first 15 participants were scanned with a Neuromag Vectorview™ MEG (Elekta Oy, Helsinki, Finland) and the other 27 with a Neuromag Triux™ MEG (MEGIN, Helsinki, Finland) due to a system upgrade. These neuromagnetometers have identical sensor layout (i.e., 102 magnetometers and 102 pairs of orthogonal planar gradiometers) and only differ in sensor dynamic range and background magnetic environment, neither of which substantially affect data quality after preprocessing. In particular, previous research mixing resting-state recordings from these two systems did not disclose any significant difference (Coquelet et al., 2020b, 2020a; Naeije et al., 2020; Sjøgård et al., 2020a, 2020b). Therefore, we did not take the MEG system type into account in later analyses.

Neuroelectric activity was measured with a MEG-compatible, 256-channel scalp EEG system (low-pass: 450 Hz; sampling frequency: 1 kHz) based on low profile, silver chloride-plated carbon-fiber electrode pellets (MicroCel Geodesic Sensor Net with Net Amp GES 400, Electrical Geodesics Inc., Magstim EGI, Eugene, Oregon, USA). The reference electrode was placed at Cz and all impedances were kept below 50 kΩ. A 100-ms long square-pulse trigger signal was generated by the MEG system electronics every second and fed to the EEG amplifier in order to enable clock synchronization of both systems. The location of the head position indicator coils, scalp EEG electrodes, and approximately 200 scalp points were determined with respect to anatomical fiducials using an electromagnetic tracker (Fastrack, Polhemus, Colchester, Vermont, USA).

Participant’s high-resolution 3D T1-weighted cerebral magnetic resonance images (MRIs) were acquired on a 1.5 T MRI scanner (Intera, Philips, The Netherlands) after the MEG/EEG recordings.

### 2.3. Data preprocessing

The MEG data were preprocessed using signal space separation (Taulu et al., 2005) to subtract environmental magnetic noise and correct for head movements (Maxfilter v2.1, Elekta Oy, Helsinki, Finland). No bad channels were detected in the process. For EEG data, we started by eliminating 84 electrodes placed on cheeks and neck as they often suffered from excessive muscle artefacts or poor skin contact, leaving 172 scalp-matched electrodes. Remnant bad channels were then automatically detected and removed using artifact subspace reconstruction (Kothe and Makeig, 2013) as implemented in EEGLAB ((Delorme and Makeig, 2004); EEGLAB v2019.0, https://sccn.ucsd.edu/eeglab/index.php) (number of bad channels: 10.6 ± 4.1 out of 172, range: 4–21). Cardiac, ocular and remaining system artifacts were further eliminated from MEG and EEG data separately, using an independent component analysis of band-passed (1–40 Hz) signals ((Vigário et al., 2000); FastICA v2.5, http://www.cis.hut.fi/projects/ica/fastica, with dimension reduction to 30 components, symmetric approach, and cubic nonlinearity contrast). Artefactual components were identified by visual inspection and regressed out of the full-rank data (number of components removed for MEG: 3.6 ± 1.1, range: 2–7; for EEG: 13.9 ± 3.3, range: 9–21). Bad EEG electrodes were subsequently reconstructed using spherical spline interpolation (Perrin et al., 1989) and EEG scalp topographies were spatially filtered (Michel and Brunet, 2019) to remove any last local outlier. The resulting EEG data were then re-referenced to the average across the 172 scalp electrodes. Finally, the synchronization of MEG and EEG signals was ensured by temporal realignment based on the trigger signal.

Separate forward models for MEG and EEG were computed based on the participants’ MRI, segmented beforehand using the FreeSurfer software (FreeSurfer v6.0; Martinos Center for Biomedical Imaging, Massachusetts, USA; https://surfer.nmr.mgh.harvard.edu, freesurfer-x86_64-linux-gnu-stable6-20170118). The coordinate systems of MEG and EEG were co-registered to the MRI coordinate system using the three anatomical fiducials for initial estimation and the head-surface points to manually refine the surface co-registration (MRIlab, MEGIN Data Analysis Package 3.4.4, MEGIN, Helsinki, Finland). The source space was built by placing three orthogonal current dipoles at each point of a grid derived from a regular 5-mm grid cropped within the Montreal Neurological Institute (MNI) template MRI volume and non-linearly deformed onto each participant’s MRI with the Statistical Parametric Mapping software (SPM12, Wellcome Centre for Neuroimaging, London, UK; https://www.fil.ion.ucl.ac.uk/spm). The forward models were then computed on this source space using the one-layer boundary element method (BEM) for MEG and the three-layer BEM with default conductivity values for EEG (as used and discussed in (Coquelet et al., 2020a)) implemented in the MNE-C suite (MNE-C v2.7.3, Martinos Center for Biomedical Imaging, Massachusetts, USA; https://mne.tools/stable/index.html). The EEG forward models were also re-referenced to their average across the 172 scalp electrodes. This source space grid and forward models were necessary for the construction of state brain maps described below.

### 2.4. Microstate clustering

Microstate inference from EEG data followed standard steps (for reviews, see, e.g., (Khanna et al., 2015; Michel et al., 2009; Michel and Koenig, 2018)) and was performed using the EEGLAB plugin for microstate analysis (v1.1, http://www.thomaskoenig.ch/index.php/software/microstates-in-eeglab) that we also adapted to MEG. Microstates were built from wideband filtered (4–30 Hz) EEG/MEG sensor signals. In the case of MEG, we focused on planar gradiometers as they disclose the highest signal-to-noise ratio (Hari and Puce, 2017) and combined each pair of orthogonal sensors using their Euclidean norm. For comparability with the power envelope signals inputted to the HMM (see below), sensor signals were here downsampled at 10 Hz using a moving-window average with 75% overlap, leading to an effective downsampling rate of 40 Hz. That said, since the microstate literature commonly uses higher sampling rates instead (Khanna et al., 2015; Michel et al., 2009; Michel and Koenig, 2018), we also applied the analysis to signals downsampled at 200 Hz (supplementary material S1).

The first step of the microstate analysis consists in a two-level clustering of time-varying sensor topographies in order to define the spatial signature of each microstate. Atomize-agglomerate hierarchical clustering (AAHC; (Tibshirani and Walther, 2005)) was used to partition each individual dataset into a number *K* of prototypal topographical maps determined so as to maximize spatial variance, a.k.a. global field power (GFP). Briefly, AAHC starts from instantaneous sensor maps and iteratively builds clusters by breaking one cluster into its constituent maps (atomization) and reassigning each of them to the cluster whose topography best fits theirs in terms of absolute spatial correlation (agglomeration). In this algorithm, the topography associated to a cluster is defined as the principal component of its constituent maps, and the cluster to atomize at each iteration is chosen deterministically as the one with least GFP. This procedure ensures that microstates are explicitly geared towards the detection of recurring patterns of highest GFP. The number *K* = 4 of clusters was fixed in accordance with the literature ((Koenig et al., 1999); for a review, see (Michel and Koenig, 2018)). The resulting set of individual-level topographies were then subjected to a full permutation procedure (Koenig et al., 1999) in order to obtain the final four group-level microstate topographies.

It is noteworthy that this two-level clustering approach is common in the microstate literature but differs from the group HMM approach (see below), so for better comparability we also considered a “group AAHC” applied to instantaneous spatial maps across all subjects at once (supplementary material S2). Also noteworthy is the fact that AAHC is restricted to time points corresponding to local maxima of the GFP time series in order to reduce computational complexity (Khanna et al., 2015). Since by design the HMM does not involve such subselection of time points, we applied microstate clustering to the unrestricted, continuous signals as well (supplementary material S3).

The second step consists in obtaining a binary time series of microstate activation/inactivation. We defined here these time series using the criterion that the microstate active at any given time point be the one whose topography best fits (again in terms of absolute spatial correlation) the instantaneous sensor topography at this time point (Brunet et al., 2011). Microstate activation is thus *exclusive*, i.e., two microstates cannot be simultaneously active, and *complete*, i.e., a microstate is active at any time. Importantly, this basic criterion is fairly close in spirit to the Viterbi algorithm used in the HMM (as explained below), but it is frequently altered in the microstate literature by using a temporally smoothed version of these binary time series (da Cruz et al., 2020; D’Croz-Baron et al., 2019; Krylova et al., 2020; Pascual-Marqui et al., 1995; Sikka et al., 2020). The Viterbi algorithm does not involve such *ad-hoc* temporal smoothing, so we primarily analyzed the raw microstate time series for better comparability with the HMM. Nevertheless, we assessed the effect of such smoothing on microstate temporal properties (see below) using the popular approach whereby microstate activation is determined as described above but at GFP peaks only and is then extended between these peaks by nearest-neighbor interpolation (Krylova et al., 2020; Sikka et al., 2020).

### 2.5. Hidden Markov modeling of power envelopes

The HMM was inferred from MEG/EEG power time courses estimated as the Hilbert envelope of wideband filtered (4–30 Hz) sensor signals. It was performed using the GLEAN toolbox (GLEAN0.3, https://github.com/OHBA-analysis/GLEAN) and adapting the pipeline described in (Baker et al., 2014), originally applied to MEG source power, for both MEG gradiometer and EEG sensor power. The focus on continuous power envelopes makes the HMM analysis geared towards the detection of power bursts (van Ede et al., 2018). Importantly, the HMM inference was done here at the sensor level for better comparability with the microstate analysis (see supplementary material S4 for the correspondence with a source-level HMM). Individual datasets of envelope signals were downsampled at 10 Hz using a moving-window average with 75% overlap (effective downsampling rate: 40 Hz), demeaned and normalized by the global variance across sensors, concatenated temporally across subjects to design a group-level analysis, and finally projected onto their *N* first principal components for dimensionality reduction prior to HMM inference (Baker et al., 2014). The dimension *N* was chosen so as to explain a comparable fraction of variance across the two modalities. Specifically, *N* = 10 components were retained for EEG and *N* = 41 for MEG, which corresponded to 81% of explained variance in both cases. This approach takes into account the intrinsic difference in spatial smoothness of MEG and EEG. (See however supplementary material S5 for a version of the EEG power envelope HMM with *N* = 41 instead, which retains more than 99% of EEG power envelope variance.)

A HMM with *K* = 6 states (Quinn et al., 2018) was then inferred from the *N* principal component time courses using variational Bayesian optimization, under several assumptions such as the normality of the observation model or the prior that hidden model parameters follow conjugate distributions (making a parametric optimization possible; for further details, see ((Rabiner, 1989; Rezek and Roberts, 2005)). Of note, the low dimensionality for EEG (*N* = 10, presumably due to high spatial smoothness of EEG; see, e.g., (Coquelet et al., 2020a)) was still sufficient to infer *K* = 6 states. The HMM optimization algorithm was run ten times, each with different initial conditions, and the model with lowest free energy was retained (Baker et al., 2014). Binary time series of most probable, temporally exclusive, and complete state activation were then derived using the Viterbi algorithm (Rezek and Roberts, 2005).

Importantly, the mere difference in number of states for the HMM and microstate analyses may trivially induce discrepancies between them. We controlled for this possibility by repeating the HMM with *K* = 4 (see supplementary material S6).

### 2.6. State temporal properties and power maps

State activation time series allowed to compute several summary statistics of the temporal behavior of microstates or HMM states, such as their mean lifetime (mean duration of state activation events) and their fractional occupancy (fraction of the total recording time during which the state is active). The global effects of recording modality and state clustering algorithm on these statistics were assessed using two-sided paired Student’s *t* tests at *p* < 0.05 applied to their average across the *K* states.

Activation time series also allowed to produce spatial maps locating where in the brain power increases or decreases occur upon state activation. The first step in this construction was to estimate the power envelope of source activity. Minimum norm estimation (Dale and Sereno, 1993) was employed as regularized inverse for projecting the wideband filtered (4–30 Hz) MEG (gradiometers only, see (Garcés et al., 2017)) and EEG signals onto the 5 mm, dipolar source grid associated to the forward models. The noise covariance matrix was estimated individually on the basis of 5 minutes of empty-room data for MEG (with signal space separation and 4–30 Hz wideband filtering), and as the identity projected in the sensor subspace corresponding to the average reference for EEG. The regularization parameter was estimated from the consistency condition derived in (Wens et al., 2015). Each three-dimensional dipole time series was projected onto the direction of maximum variance, and the Hilbert envelope of the resulting source signal was then extracted and downsampled as described above for the sensor signals.

Brain power maps were then obtained by computing the partial correlation between each state activation time series and these source power envelope signals concatenated across subjects (Baker et al., 2014). Corresponding maps could also be derived at the individual level by mere restriction of these partial correlations within each subject. In the HMM case, this procedure was also performed at the level of sensor power envelope signals used for state inference. All these maps were thresholded statistically using two-tailed parametric correlation tests at *p* < 0.05 against the null hypothesis that Fisher-transformed correlations follow a Gaussian with mean zero and SD 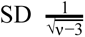, where ν = *N_tdof_* − (*K* − 1) . The number *N_tdof_* of temporal degrees of freedom was estimated as one-quarter of the total number of time samples in group-concatenated envelope signals at 40 Hz sampling frequency, to take into account the 75% overlap in the envelope downsampling. The subtraction of *K* – 1 degrees of freedom is due to the regression inherent to the partial correlation. The critical *p*-value was Bonferroni corrected with the number of independent states (i.e., *K* – 1) multiplied by the number of spatial degrees of freedom estimated from the rank of the forward model (Wens et al., 2015), i.e., 58 for MEG and 32 for EEG. Statistical thresholding on state power maps was thus slightly tighter for MEG than EEG, which is merely a reflection of the higher spatial smoothness in EEG data (Coquelet et al., 2020a).

### 2.7. State correlation analyses

The spatial and temporal profiles of each pair of states were compared quantitatively using correlation analyses, in order to assess the effect of the recording modality (MEG vs. EEG) and of the state clustering model (microstates vs. HMM). The spatial similarity of two states was assessed using Pearson correlation of their source-level brain power maps, and their tendency to co-activate using Spearman correlation of their binary activation time series, both computed within each subject. Statistical significance was then established using one-sided one-sample parametric *t*-tests against the null hypothesis that the group-averaged sample correlation vanishes (reflecting the absence of topographical resemblance or of temporal co-activation between two states) and with the alternative hypothesis that this average is positive (reflecting significant topographical overlap or temporal co-activation). The significance level was set to *p* < 0.05 Bonferroni corrected for the number of possible state pairs included in the comparison at stake.

### 2.8. Data and code availability statement

The MEG/EEG data and analysis code used in this study will be made available upon reasonable request to the corresponding author.

## 3. Results

### 3.1. Microstates

Figure 1 shows the spatial signature of the four microstates derived from EEG and MEG resting-state data (AAHC with *K* = 4 applied to 40 Hz-downsampled sensor maps at time points of local GFP maxima).

**Figure 1:**
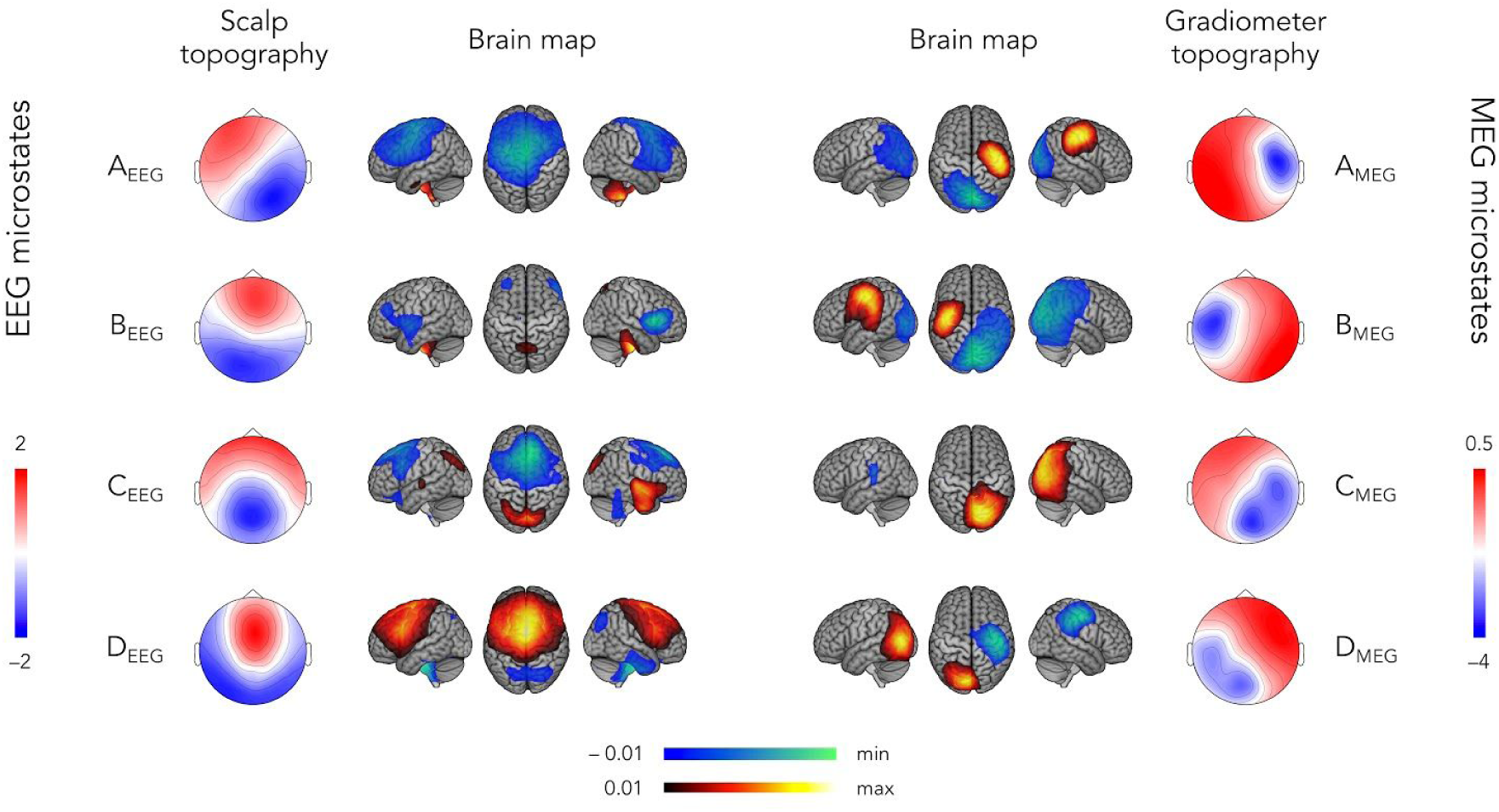
Spatial signature of EEG (**left**) and MEG (**right**) microstates. The scalp topography of EEG microstates (four-cluster AAHC of the 40 Hz-downsampled sensor maps at time points of local GFP maxima) is shown on the far left and the corresponding brain maps on the middle left. The gradiometer topography of MEG microstates is shown on the far right and the corresponding brain maps on the middle right. The scales for sensor topographies correspond to standardized *z* scores. Positive (negative) values in the brain maps indicate increasing (decreasing) power upon microstate activation. These maps are thresholded statistically and the lower/upper scales are adapted to the minimum/maximum values.

The EEG microstate analysis allowed to reproduce canonical scalp topographies (Fig. 1, left) well established in the literature (Michel and Koenig, 2018). Each microstate displayed a scalp potential distribution reminiscent of one current dipole (notwithstanding the difficulty of interpreting scalp EEG in this way; see, e.g., (Hari and Puce, 2017)) approximately located centrally and oriented along the left frontal to right posterior line (microstate A_EEG_), the frontal to posterior midline (microstates B_EEG_ and C_EEG_, with somewhat more anterior location or more inferior frontal orientation for the latter), or predominantly vertically (microstate D_EEG_). The corresponding brain maps shown in Fig. 1 (left) identify the sources exhibiting significant power increase (positive values) or decrease (negative values) upon microstate activation. Microstates A_EEG_, C_EEG_, and D_EEG_ were dominated by a power modulation peaking at a midline frontal source, confirming the prominently dipolar nature of their scalp topography. Specifically, microstate A_EEG_ was characterized by a power decrease at this source location. Microstate C_EEG_ exhibited a similar (although slightly more anterior) power decrease but also power increases at the precuneus and fronto-temporal areas. Microstate D_EEG_ involved an opposite pattern, i.e., power increase at the midline frontal source and power decrease at the precuneus. On the other hand, microstate B_EEG_ was characterized by power decreases at bilateral inferior frontal sources and a power increase at the precuneus. It is noteworthy that its scalp topography is indeed compatible with two bilateral temporal current dipoles. All four brain power maps also disclosed deep cerebellar patterns that may be related to EEG source reconstruction errors.

Although the spatial signature of these microstates matches the literature, their temporal statistics differed substantially, with mean lifetimes shorter than expected (mean ± SD: 37 ± 2 ms, range: 35–38 ms; see Table 1, left). These lifetimes were even shorter (mean ± SD: 14 ± 1 ms, range: 13–15 ms) when clustering EEG topographies at a higher sampling rate (200 Hz; see also supplementary material S1). This discrepancy was merely imputable to the absence of temporal smoothing on the microstate activation time series, as the interpolation approach allowed to recover typical lifetimes (mean ± SD: 126 ± 8 ms, range: 121–138 ms). Fractional occupancies ranged from 19% to 27% (Table 1, left) and were not substantially affected by temporal smoothing (23–28%).

**Table 1:** Mean lifetimes and fractional occupancies (mean ± SD) associated with each microstate inferred from EEG or MEG topographies at 40 Hz sampling rate and without temporal smoothing on microstate activation time series.

| EEG microstates |  |  | MEG microstates |  |  |
| --- | --- | --- | --- | --- | --- |
|  | Mean lifetimes (ms) | Fractional occupancies (%) |  | Mean lifetimes (ms) | Fractional occupancies (%) |
| $A_{\text{EEG}}$ | $38 \pm 3$ | $26.9 \pm 4.4$ | $A_{\text{MEG}}$ | $32 \pm 1$ | $14.7 \pm 1.8$ |
| $B_{\text{EEG}}$ | $35 \pm 2$ | $19.5 \pm 3.9$ | $B_{\text{MEG}}$ | $37 \pm 2$ | $25.1 \pm 3.7$ |
| $C_{\text{EEG}}$ | $37 \pm 3$ | $25.8 \pm 4.8$ | $C_{\text{MEG}}$ | $47 \pm 2$ | $41.1 \pm 2.9$ |
| $D_{\text{EEG}}$ | $38 \pm 2$ | $27.6 \pm 4.3$ | $D_{\text{MEG}}$ | $34 \pm 2$ | $19.1 \pm 2.3$ |

A similar analysis applied to MEG gradiometer signals revealed microstates that were also dominated by dipolar sensor topographies (Fig. 1, right). Microstates A_MEG_ and B_MEG_ were characterized by gradiometers peaking respectively above the right and the left parietal sensors, which was explained by unilateral power increases at the sensorimotor cortices. These two microstates also involved an occipital power decrease. Microstates C_MEG_ and D_MEG_ disclosed respectively right and left parieto-occipital gradiometer activity corresponding to unilateral occipital power increases. The brain map for microstate D_MEG_ also showed right sensorimotor power decrease. The neural generators behind MEG microstates thus appeared qualitatively different from those of EEG microstates. On the other hand, their mean lifetimes (mean ± SD: 37 ± 7 ms, range: 32–47 ms; Table 1, right) were similar (*t*_41_ = 1.8, *p* = 0.08). Fractional occupancies appeared less homogenous for MEG (14–41%; Table 1, right) than for EEG, with microstate C_MEG_ showing the highest fractional occupancy. Analogously to the EEG case, temporal smoothing lengthened MEG microstates lifetimes (mean ± SD: 114 ± 21 ms, range: 94–141 ms) but did not affect fractional occupancies (17–38%), and increasing the signal sampling rate at 200 Hz further shortened lifetimes (mean ± SD: 11 ± 3 ms, range: 8–16 ms; see also supplementary material S1).

Several variations of the microstate clustering approach are explored in supplementary materials. These analyses show that the above features of microstates are robust against methodological changes such as increasing the signal sampling rate (except for the effect on lifetimes, see supplementary material S1), using group clustering (supplementary material S2) or lifting the restriction to GFP local maxima (supplementary material S3).

### 3.2. Hidden Markov model states

Figure 2 depicts the spatial signature of the six HMM states inferred from resting-state MEG and EEG sensor-level power envelopes. States were sorted and labeled in order to pair EEG states (Fig. 2, left) and MEG states (Fig. 2, right) with the best apparent spatial correspondence. An important difference with Fig. 1 is that sensor maps in Fig. 2 locate power increases or decreases upon state activation, analogously to the brain maps. As a matter of fact, both sensor and brain maps are directly comparable for MEG because planar gradiometers are sensitive to source activity just beneath them (Hari and Puce, 2017). This comparison is less straightforward for EEG. For this reason, we mostly focus on a description of their brain maps.

**Figure 2:**
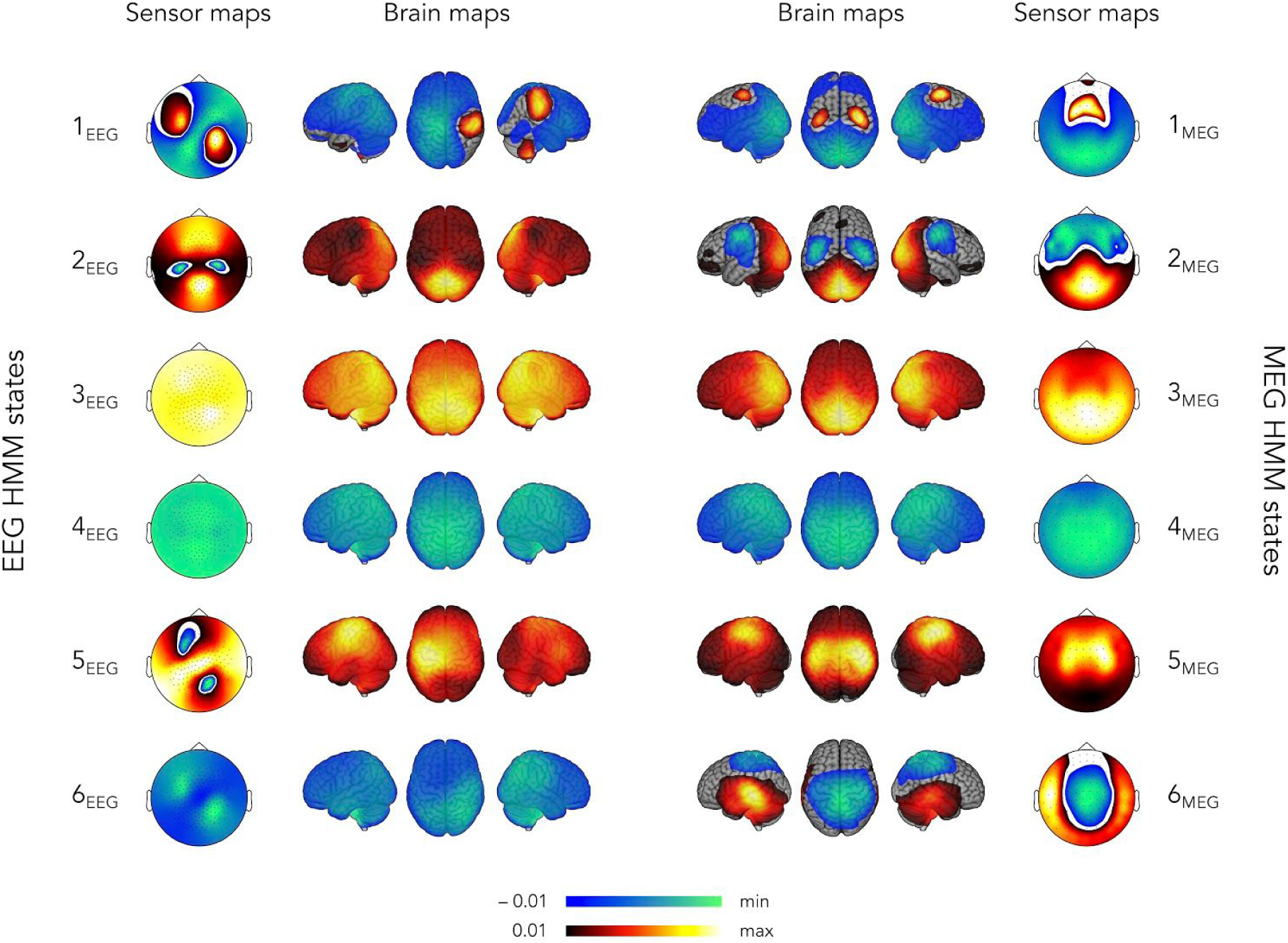
Spatial signature of EEG (**left**) and MEG (**right**) sensor-level power envelope HMM states. Both sensor and brain maps locate power increases (positive values) and decreases (negative values) upon state activation. These maps are thresholded statistically and the lower/upper scales are adapted to the minimum/maximum values. States were paired based on the visual correspondence of the brain maps.

We start with the MEG states (Fig. 2, right) as this is the most conventional application of the power envelope HMM. In accordance with previous studies (Baker et al., 2014; Brookes et al., 2018; Coquelet et al., 2020b), these states identified power modulations within well-known intrinsic functional networks, here the sensorimotor network (SMN), the visual occipital network (VoN), the posterior part of the default-mode network (pDMN; encompassing the precuneus), and a presumed auditory network (AN). More specifically, state 1_MEG_ involved SMN power activation along with pDMN power deactivation, while state 2_MEG_ displayed an opposite pattern of SMN deactivation and VoN activation. These two states are thus reminiscent of a dynamic competition between the SMN and pDMN/VoN (Wens et al., 2019). States 3_MEG_ and 4_MEG_ identified pDMN activation and deactivation, respectively. Of note, similar states involving precuneus activity were also identified in previous works (Coquelet et al., 2020b; Puttaert et al., 2020) and not in others (e.g., (Baker et al., 2014; Brookes et al., 2018)) due to different choices of source projection (for details, see (Sjøgård et al., 2019)). State 5_MEG_ corresponded to the activation of the SMN in isolation (rather than in competition with the pDMN/VoN, as in states 1_MEG_ and 2_MEG_). Finally, state 6_MEG_ involved AN activation alongside power deactivation at the precuneus, which is once again reminiscent of a dynamic cross-network competition (Wens et al., 2019). Mean lifetimes were significantly longer than for the non-smoothed MEG microstates (mean ± SD: 151 ± 31 ms; range: 128–211 ms; *t*_41_ = 31.6, *p* = 0), and fractional occupancies ranged between 7% and 23% (Table 2, right).

**Table 2:** Mean lifetimes and fractional occupancies (mean ± SD) associated with each of the six HMM states inferred from EEG or MEG power envelope signals.

| EEG HMM |  |  | MEG HMM |  |  |
| --- | --- | --- | --- | --- | --- |
|  | Mean lifetimes (ms) | Fractional occupancies (%) |  | Mean lifetimes (ms) | Fractional occupancies (%) |
| State 1<sub>EEG</sub> | 136 ± 19 | 17 ± 3.7 | State 1<sub>MEG</sub> | 138 ± 34 | 22.2 ± 10.9 |
| State 2<sub>EEG</sub> | 145 ± 39 | 13.3 ± 4.5 | State 2<sub>MEG</sub> | 144 ± 41 | 16.4 ± 8.2 |
| State 3<sub>EEG</sub> | 204 ± 70 | 8.4 ± 1.4 | State 3<sub>MEG</sub> | 153 ± 52 | 7.5 ± 3.2 |
| State 4<sub>EEG</sub> | 226 ± 109 | 22.4 ± 9 | State 4<sub>MEG</sub> | 211 ± 98 | 23.1 ± 10.6 |
| State 5<sub>EEG</sub> | 137 ± 20 | 14.6 ± 4.7 | State 5<sub>MEG</sub> | 128 ± 40 | 12.5 ± 7.7 |
| State 6<sub>EEG</sub> | 141 ± 23 | 24.3 ± 6.2 | State 6<sub>MEG</sub> | 130 ± 54 | 18.3 ± 15.4 |

The HMM states inferred from EEG involved power modulations within intrinsic networks similar to the MEG states, although not with the same degree of bilaterality (Fig. 2, left). State 1_EEG_ was characterized by the activation of the right part of the SMN alongside a power decrease in the left precuneus, and as such may be viewed as a unilateral version of MEG state 1_MEG_. This state was the only EEG state exhibiting both power increases and decreases. The VoN activation state 2_EEG_ was comparable to state 2_MEG_ but lacked SMN deactivation, and the pDMN states 3_EEG_ and 4_EEG_ closely matched states 3_MEG_ and 4_MEG_. State 5_EEG_ was characterized by a power increase in the left part of the SMN, so it appeared as a unilateral version of state 5_MEG_. Finally, state 6_EEG_ consisted in a right-hemispheric posterior parietal power decrease, which was thus qualitatively different from the AN/precuneus state 6_MEG_. The mean lifetime of these states (mean ± SD: 165 ± 40 ms, range: 136–204 ms; see Table 2, left) was significantly longer than the non-smoothed EEG microstates (*t*_41_ = 31.46, *p* = 0) and the MEG HMM states (*t*_41_ = 4.83, *p* = 1.9 × 10^−5^). Fractional occupancies were between 8% and 24%, which is also similar to those observed using MEG (Table 2).

Interestingly, the brain maps shown in Fig. 2 and reconstructed from sensor-level HMM states were qualitatively similar to those obtained from HMM states directly inferred from source power envelopes (supplementary material S4). For EEG, increasing the dimensionality of the data inputted to the HMM algorithm to the same dimension used in MEG, also led to qualitatively similar states, although with a higher degree of bilaterality for some states (supplementary material S5). Finally, reducing the number of states to four led to HMM states closely related to states 2_MEG_–4_MEG_ for MEG and to states 1_EEG_, 3_EEG_–5_EEG_ for EEG (supplementary material S6).

### 3.3. State correlations

Figure 3 details the group-level spatial (Fig. 3, top) and temporal (Fig. 3, bottom) correlations among states, the former to quantify the spatial correspondence discussed qualitatively above and the latter, their tendency to co-activate. The two left columns assess the effect of recording modality (MEG vs. EEG) on microstates (Fig. 3, first column) and HMM states (Fig. 3, second column) and the two right columns, the effect of state clustering algorithm (microstate vs. HMM) for both EEG (Fig. 3, third column) and MEG (Fig. 3, fourth column). Of note, these temporal correlations were obtained using non-smoothed microstate activation time series, as no temporal smoothing is explicitly applied to HMM state activation time series.

**Figure 3:**
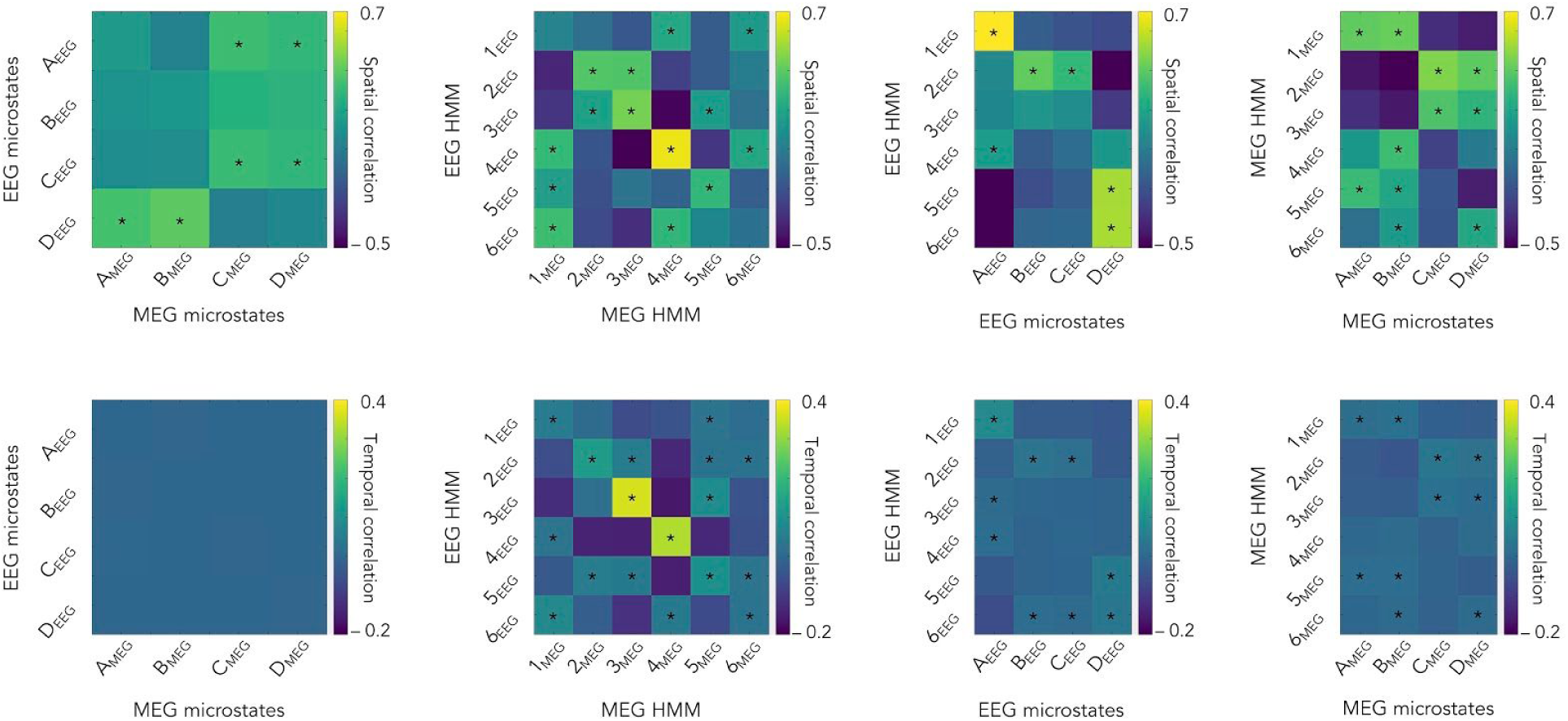
Spatial (**top**) and temporal (**bottom**) state correlations. Each matrix shows the group-level correlation values comparing: EEG microstates vs. MEG microstates (**first column**; corresponding to four-cluster AAHC of the 40 Hz-downsampled sensor maps at time points of local GFP maxima), EEG HMM states vs. MEG HMM states (**second column**; six-state HMM of sensor-level power envelopes), EEG HMM states vs. EEG microstates (**third column**), and MEG HMM states vs. MEG microstates (**fourth column**). Temporal correlations were obtained from the raw (non-smoothed) microstate activation time series. The same correlation scale is used across the four comparisons. Stars denote significant correlations after Bonferroni correction for the number of state pairs involved in each comparison.

The comparison of EEG vs. MEG microstates confirmed the absence of a clear relationship. Some cross-modal pairs did disclose significant spatial correlations (Fig. 3, top of left column; significant *R* > 0.09, *t*_41_ > 3.21, *p* < 0.021 Bonferroni corrected for 16 comparisons). They could be explained by a gross overlap of their power maps presumably due to their intrinsic blurriness, i.e., EEG microstates A_EEG_ and C_EEG_ tended to exhibit posterior power increases and antero-central power decreases as did the MEG microstates C_MEG_ and D_MEG_, and reversely for microstates D_EEG_, A_MEG_, and B_MEG_ (Fig. 1). More importantly, the corresponding temporal correlations were not significant with very small effect sizes (Fig. 3, bottom of left column; *R* < 0.002, *t*_41_ < 1.14, *p* > 0.13 uncorrected), indicating that EEG and MEG microstates scarcely co-activated at all.

On the other hand, Fig. 3 (second column) revealed a number of significant correlations between MEG and EEG HMM states, both spatially (significant *R* > 0.15, *t*_41_ > 3.85, *p* < 7.2 × 10^−3^ Bonferroni corrected for 36 comparisons) and temporally (significant *R* > 0.025, *t*_41_ > 4.02, *p* < 4.2 × 10^−3^ corrected), and with higher effect sizes and smaller *p* values than microstates. The qualitative pairing of MEG and EEG states based on their maps (Fig. 2) was reflected in significance along the diagonal of the correlation matrix (Fig. 3, top of second column), with particularly high effect size and low *p* value for the pDMN deactivation state 4_MEG_/4_EEG_ (*t*_41_ = 32.73, *p* = 0). The two exceptions were states 1_MEG_/1_EEG_ (where the correlation did not reach significance presumably due to the sign reversal above the left sensorimotor cortex) and states 6_MEG_ and 6_EEG_. Analogously to the case of microstates, off-diagonal significance may be a reflection of spatial blurriness. Temporal correlations followed a similar pattern (Fig. 3, bottom of second column), with the pDMN states 3_MEG_/3_EEG_ and 4_MEG_/4_EEG_ standing out regarding their effect size and *p* value (*t*_41_ > 14.11, *p* = 0). Globally, the HMM inference on power envelopes was thus able to identify common states across the two recording modalities.

We turn now to the comparison of microstates and HMM states within each modality. Spatial correlations appeared significant in a number of microstate/HMM state pairs (EEG: Fig. 3, top of third column, significant *R* > 0.16, *t*_41_ > 3.2, *p* < 0.031 Bonferroni corrected for 24 comparisons; MEG: Fig. 3, top of fourth column, significant *R* > 0.15, *t*_41_ > 4.02, *p* < 2.29 × 10^−3^ corrected). For EEG, these correlations appeared to mainly reflect spatial blurriness since microstate and HMM states power maps peaked at distinct locations, except for similar power increases at the visual cortices (microstates B_EEG_, C_EEG_ and HMM state 2_EEG_, see Figs. 1 and 2, left). Spatial similarities among MEG states exhibited higher effect sizes and lower *p* value. Accordingly, Figs. 1 and 2 (right) indicated some degree of co-localization in microstate and HMM state power modulation within the SMN (activation for microstates A_MEG_, B_MEG_ and HMM states 1_MEG_, 5_MEG_; deactivation for microstate D_MEG_ and HMM state 2_MEG_) and VoN (activation for microstates C_MEG_, D_MEG_ and HMM states 2_MEG_, 3_MEG_). Temporal correlations followed once again a somewhat similar pattern of significance (EEG: Fig. 3, bottom of third column, significant *R* > 0.01, *t*_41_ > 3.19, *p* < 0.034 corrected; MEG: Fig. 3, bottom of fourth column, significant *R* > 0.007, *t*_41_ > 3.52, *p* < 0.013 corrected). However, the raw value of these significant correlations remained low (EEG: *R* < 0.09, MEG: *R* < 0.04), so temporal co-activations were marginal.

## 4. Discussion

This study used simultaneous MEG/EEG recordings at rest to compare two notions of discrete metastable brain states, i.e., microstates and power envelope HMM states. We found that microstates were not reproducible across the two recording modalities, i.e., microstates inferred from MEG signals did not correspond to the canonical EEG microstates. On the other hand, MEG and EEG HMM states identified transient activations of the same intrinsic functional networks, although with a limited temporal correspondence. We also found no evidence that EEG microstates and HMM states share common neural dynamics. In fact, contrary to our expectation based on the literature (Baker et al., 2014; Michel and Koenig, 2018), all microstates were substantially less stable in time than the HMM states. That said, the MEG version of microstates involved power activity within the same networks as HMM states, but was restricted to isolated nodes of these networks, and with a poor temporal correspondence.

### 4.1. Microstates and power envelope HMM states probe different aspects of electrophysiological power bursts

The primary result of this paper is that microstates and power envelope HMM states differ substantially, both in the localization of the brain areas they (de)activate and in their temporal stability. These two state clustering algorithms share the common goal of identifying patterns of high-power electrophysiological activity that repeat at rest, so this raises the questions of what methodological features lead to this discrepancy, and what aspect of brain functional dynamics they are preferentially sensitive to. The fundamental distinction discussed here is that (i) microstates focus on high-power activity by biasing the topographical clustering to time points of locally maximum GFP (Michel and Koenig, 2018), whereas (ii) the power envelope HMM encodes states based on the spatial patterns of continuous-time oscillatory power (Baker et al., 2014).

The GFP maximization for microstate topographies is fully built-in in the AAHC algorithm (Murray et al., 2008). In fact, the convergence of the AAHC with and without explicit restriction to GFP peaks indicates that microstates are mostly specific to time points of locally maximal GFP. This concurs with the reportedly high levels of EEG topographical dissimilarities in between GFP peaks (Skrandies, 1990) and with the difficulty of discrete microstates to model continuous EEG recordings (Mishra et al., 2020). Accordingly, in our data, the duration of microstate activation appeared very short, and was actually only slightly above the minimum timescale allowed by signal processing (at least in the absence of temporal smoothing). A mean lifetime of 37 ms is only 150% the 25 ms timestep of our signals sampled at 40 Hz, and the fact that it decreased by merely increasing the sampling rate indicates that microstates are actually even shorter lived. Classical lifetimes of 120 ms appear to require an *ad-hoc* temporal interpolation procedure that does not reflect the raw GFP peak events underlying microstate clustering nor the high topographical dissimilarities in between these events (Skrandies, 1990). Extrapolating the observation that raw microstate lifetimes are 150% the timestep would have led us to expect a mean lifetime of about 7 ms at 200 Hz sampling rate (corresponding to a 5 ms timestep), but our data proved it twice longer. This is presumably a sign that microstates do reflect neural events, since neurophysiological activity as recorded by MEG/EEG should typically not occur over timescales shorter than the 10 ms duration of postsynaptic potentials (Baillet, 2017; Buzsáki et al., 2012), whereas pure noise events can be as short as the timestep. Microstates thus appear to probe quasi-instantaneous electrophysiological events.

Further understanding what these microstate events represent requires careful consideration of the notion of GFP. Instantaneous GFP (spatial variance of time-dependent sensor topographies) is not trivially synonymous with instantaneous global power (magnitude squared signal summed over all sensors). For EEG, the two concepts coincide only when using the average reference (where the potential summed over all electrodes is constrained to vanish), which approaches the idealized reference to infinity because asymptotically vanishing electric potentials generated by current dipoles inside the brain integrate to zero over the scalp (Bertrand et al., 1985), at least to some approximation (Yao, 2017). From a physical perspective, GFP maximization of EEG microstates is thus theoretically equivalent to global power maximization. In practice though, the GFP formulation is preferred because it is strictly independent of the choice of reference (Murray et al., 2008; Skrandies, 1990). No such subtlety arises with MEG, where GFP and global power coincide because neuromagnetic field patterns generated by dipolar brain sources also sum up approximately to zero over whole-head-covering sensor arrays (meaning in a sense that the “average reference” holds automatically for MEG). Thus, the quasi-instantaneous microstate events correspond to moments of high global power. Given that spontaneous electrophysiological activity exhibits power bursts (Hari and Salmelin, 1997; van Ede et al., 2018), microstates may be expected to probe short-time events of maximum power within power bursts. More specifically, since the EEG/MEG spectrum is dominated by the alpha band (Hari and Puce, 2017), microstates are bound to be driven by, and phase-locked to, moments of high-amplitude alpha rhythms within alpha bursts (von Wegner et al., 2021).

In any case, by focusing on quasi-instantaneous and temporally discrete electrophysiological events of high power, microstates provide at best a partial characterization of power bursts. Their full exploration requires instead to focus on the transitioning between low and high power. By running over the whole power envelope signal, the HMM is more sensitive to such transitions and may thus be better suited to fully capture bursting activity (van Ede et al., 2018). The Markovian character of the HMM (i.e., the probability of state activation at the next time step depends on what state is currently active; see, e.g., (Rabiner, 1989)) also enforces a degree of deterministic causality that further helps detecting transient periods of sustained power burst, rather than quasi-instantaneous events of high power. Accordingly, bursts generated by brain rhythms typically last for a few hundreds of milliseconds (Hari and Salmelin, 1997), which is consistent with the typical power envelope HMM mean lifetime of 100–200 ms. The fact that these lifetimes are well above the minimum timestep allowed in our power envelope signals (in our case, 25 ms) further shows that they provide a reliable estimate of the duration of the underlying power bursts.

### 4.2. Microstates identify local neural events whereas power envelope HMM states encompass network-level activity

Besides temporal stability, microstates and power envelope HMM states also differed in their spatial distribution, with microstates exhibiting isolated power modulations and HMM states, distributed (de)activations at the network level. The first methodological reason to consider is that microstate clustering relies on topographical similarity (Michel and Koenig, 2018) whereas the HMM encodes the whole covariance structure (Baker et al., 2014; Woolrich et al., 2013). In theory, HMM states are thus driven by a mixture of power topography and intrinsic functional connectivity. This being said, the contribution of functional connectivity (more specifically, the cross-covariance feature in the HMM) may not dominate HMM state inference in practice (Vidaurre et al., 2018). In fact, functional networks can also be identified successfully using classification schemes that do not encode explicitly for envelope cross-covariance, such as the independent component analysis of power envelopes (which is, however, generally applied at slower timescales around 1 s; see, e.g., (Brookes et al., 2011; Wens et al., 2014)).

The involvement of functional networks in HMM states and the lack thereof in microstates might alternatively be rooted in their difference in temporal stability discussed at length above. The physiological process of binding distant neural populations into a functional network entails a hierarchy of timescales, from hundreds of milliseconds accessible to the HMM for certain networks (i.e., SMN, DMN and visual network) to several seconds for others (e.g., the fronto-parietal network) (Baker et al., 2014; Vidaurre et al., 2018). With lifetimes below these timescales and associated with quasi-instantaneous MEG/EEG events, microstates may thus be mostly sensitive to highly transient neural activity taking place locally without enough time to establish network-level coordination. A somewhat related hypothesis was put forth when comparing EEG microstates to fMRI networks (Britz et al., 2010; Musso et al., 2010; Yuan et al., 2012). This is also in line with our observation that MEG microstates appeared as unilateral versions of some HMM states. For example, correlation results suggested that HMM state 1_MEG_ may be viewed as a combination of microstates A_MEG_ and B_MEG_.

Closely related to this timescale argument, the spatial locality of microstates may also be viewed in light of their being phase locked to alpha rhythms (von Wegner et al., 2021). A putative “network-level” microstate involving distinct brain regions would then imply the existence of a zero-phase lag synchronization among them, and as such it would presumably not reflect neurophysiological activity. This is because zero-lag synchronization among separated brain areas evidences instantaneous interactions, which are generally thought to be non-physiological (Schoffelen and Gross, 2009; Wens, 2015) (see, however, (Sjøgård et al., 2019), for a spontaneous, nearly zero-lag correlation within the default-mode network). One way to further investigate the relationship between microstates and synchronization would be to compare them to another implementation of the HMM (Vidaurre et al., 2018) that is not applied on MEG/EEG power envelopes but on the MEG/EEG signals with time-delay embedding (Kantz and Schreiber, 2003; Takens, 1981), which gives access to classification features closely related to phase synchrony (Stam and van Dijk, 2002). Compared to the power envelope HMM, this time-embedded HMM exhibits shorter lifetimes (50–100 ms), richer spectral details, and network-level phase locking (Vidaurre et al., 2018). Given that these lifetimes are still well above the smallest accessible timestep (4 ms at the 250 Hz sampling rate used in (Vidaurre et al., 2018)) and thus cannot be deemed quasi-instantaneous, and that this network synchrony occurred at non-zero phase lag, we surmise that the time-embedded HMM provides yet another state description, more stable than microstates but more transient than power envelope HMM states. Still, it would be useful to perform such comparisons explicitly in the future.

In sum, the above considerations suggest that microstates and HMM states are sensitive to neural events occurring at different temporal and spatial scales, highly transient and isolated for the former, more stable and distributed over intrinsic networks for the latter.

### 4.3. Cross-modal comparisons reveal poor correspondence of state activations

The conclusion that microstate classification depends on highly transient events is also key to understanding the lack of qualitative correspondence between EEG and MEG microstates. This discordance contrasted with the HMM, which disclosed good spatial similarity across the two recording modalities. The pDMN state pairs 3_MEG_/3_EEG_ and 4_MEG_/4_EEG_ exhibited a substantial overlap of their activation periods, but the others lacked such strong temporal correspondence. The pDMN states were also the most stable (Coquelet et al., 2020b; Puttaert et al., 2020), suggesting that state co-occurrence rate increases with state stability. The difficulty of short-lived states to co-activate explains in particular the poor cross-modal temporal correlation for the quasi-instantaneous microstates. This observation is also in line with a previous comparison of MEG and EEG intrinsic functional connectomes, which were spatially similar but with rather discordant temporal dynamics (Coquelet et al., 2020a). The hypothesis raised to explain this result was that MEG and EEG are sensitive to different components of transient functional integration processes, but that these differences smooth out after minute-scale time averaging. Our results suggest that this smoothing effect extends to the finer timescales accessible to MEG/EEG state analyses. In fact, it fits well with the observation discussed above that some relatively stable, network-level HMM states break into highly transient, spatially local MEG microstates.

The spatial discordance between EEG and MEG microstates can then be understood on this basis. A sensitivity of EEG and MEG to different transient neural events (as hypothesized above) would lead to different GFP maxima and thus, to microstates inferred from totally different time points. More generally, the concept of GFP maxima turns out to be modality specific since it depends on the type of sensors used (here, EEG electrodes vs. MEG gradiometers). We focused in this paper on MEG microstates derived from gradiometers, but it is noteworthy that microstates based on magnetometers also poorly correlate with gradiometric microstates (data not shown). We conclude that microstates obtained with different electrophysiological modalities probe neural events occurring at different times and are thus not directly comparable. As emphasized above, HMM state inference does not depend on a modality-specific selection of time periods, which explains their better cross-modal concordance. Other potential sources of differences are the distinct sensitivity profiles of EEG and MEG, especially to purely radial dipolar sources (Hari and Puce, 2017), and the higher regional variability of sensor-brain distance with MEG arrays than with scalp EEG (Coquelet et al., 2020a). The latter impacts substantially MEG functional connectivity estimation in frontal regions from which MEG sensors are farthest (Coquelet et al., 2020a), but interestingly no such issue was clearly observable in brain maps of sub-second HMM states. Still, these differences might partially account for their poor temporal correspondence.

### 4.4. Power envelope HMM states can be inferred directly from sensor-level signals

One side result noteworthy of mention is that the HMM of sensor power signals leads to network-level states similar to the HMM of reconstructed source power considered in the seminal paper of (Baker et al., 2014) and subsequent MEG studies (Brookes et al., 2018; Coquelet et al., 2020b; Puttaert et al., 2020; Quinn et al., 2018; Sitnikova et al., 2018; Van Schependom et al., 2019; Vidaurre et al., 2018). The HMM of electrophysiological signals can thus be performed in a computationally less cumbersome way than previously done, for similar results. This might widen the perspectives of applications of the HMM-based analyses of MEG/EEG data, particularly when studying infants or patients where MRI acquisition might not be possible. Methodologically, this also frees the HMM state inference *per se* from ambiguities related to the choice of forward model (for further discussion of this aspect, see, e.g., (Coquelet et al., 2020a)) and source reconstruction algorithm. Only the imaging of state brain maps would depend on these choices. This is particularly interesting with regard to the contribution of precuneus activity to HMM state dynamics, as it was identified from MEG source power HMMs when using minimum norm estimation (Coquelet et al., 2020b; Puttaert et al., 2020) but not when using a beamformer (Baker et al., 2014; Brookes et al., 2018; Vidaurre et al., 2018) due to a suppression effect (Sjøgård et al., 2019). The states 1_MEG_, 3_MEG_/3_EEG_ and 4_MEG_/4_EEG_ obtained in this study show that sensor-level HMM is sensitive to precuneus activity, independently of source reconstruction biases. This being said, it would be interesting in the future to extend our comparative study to other implementations of the power envelope HMM, e.g., restricted to parcellated source reconstruction and with multivariate signal leakage correction (Brookes et al., 2018; Colclough et al., 2015; Sitnikova et al., 2018), and examine whether such processing steps improve the robustness of HMM state inference.

One last aspect to emphasize in the case of EEG is that, strictly speaking, the power envelope HMM is ill-defined because it relies on the concept of EEG signal power, which depends on the choice of reference. As discussed above, we focused here on the average reference, which approximates the physically ideal reference at infinity and thus presumably mitigates this issue in practice. This is in line with our observation that source-projected brain maps of sensor-level HMM states correspond to maps of source-level HMM states, the latter being based on current dipole estimates that are independent of the reference (of course, the choice of recording reference does matter, as it impacts measurement quality (Hari and Puce, 2017)). Still, sensor-level HMM state inference may be improved by using, e.g., the reference electrode standardization technique that aims at simulating a virtual reference at infinity (Yao, 2001).

### 4.5. Conclusion

This study revealed that microstates and HMM states reflect neural dynamical events probing power bursts at different timescales. The spatio-temporally local character of microstates explains their specificity to the electrophysiological recording modality at hand. For EEG, microstate analysis and the power envelope HMM appear to bring complementary information about transient neural dynamics, so we suggest that the two approaches should be considered together. On the other hand, the added value of MEG microstates may be more limited as they merely identify a short-time splitting of network-level HMM states. Both approaches allow to model fast, spontaneous bursts of electrophysiological activity occurring at sub-second timescales. As such, they represent important tools to further explore the dynamical functional architecture of the human brain.

## Acknowledgments

This study was supported by the Action de Recherche Concertée Consolidation (ARCC, “Characterizing the spatio-temporal dynamics and the electrophysiological basis of resting state networks”, ULB, Brussels, Belgium), the Fonds Erasme (Research Convention “Les Voies du Savoir”, Brussels, Belgium) and the Fonds de la Recherche Scientifique (Research Conventions: T.0109.13, Excellence of Science EOS MEMODYN (30446199); FRS-FNRS, Brussels, Belgium). Nicolas Coquelet has been supported by the ARCC and by the Fonds Erasme (Research Convention “Les Voies du Savoir”, Brussels, Belgium), and is now supported by the FRS-FNRS (Research Conventions: Excellence of Science EOS MEMODYN (30446199); Brussels, Belgium). Xavier De Tiège is Postdoctorate Clinical Master Specialist at the FRS-FNRS. Lillia Roshchupkina is FRS-FNRS Research Fellow and was previously supported by a ULB Mini-ARC grant. Mark W. Woolrich is supported by the NIHR Oxford Health Biomedical Research Center, the Wellcome Trust (098369/Z/12/Z, 106183/Z/14/Z, 215573/Z/19/Z), and the New Therapeutics in Alzheimer’s Diseases (NTAD) study supported by the UK MRC and the Dementia Platform UK. The MEG project at the CUB Hôpital Erasme is financially supported by the Fonds Erasme (Research Convention: “Les Voies du Savoir”, Brussels, Belgium). The high-density EEG project at the CUB Hôpital Erasme has been financially supported by the CUB Hôpital Erasme (Medical Council Research Grant) and by the FRS-FNRS. The authors would like to thank Maribel Pulgarin-Montoya for her help in parts of the MEG/EEG recordings.

## Author contributions

N.C., X.D.T. and V.W. designed study; N.C., L.R., X.D.T and V.W. acquired data; N.C. and V.W. contributed to analysis tools; N.C., X.D.T. and V.W. analysed data; N.C., X.D.T., L.R., P.P., S.G., M.W. and V.W. wrote and reviewed the manuscript.

## Supplementary material

### S1. Effect of increased sampling rate on microstates

The EEG microstate literature commonly relies on sampling rates higher than 40 Hz, on which we focused in the main text for consistency with the standard HMM analysis. We repeat here the main microstate analysis (*K* = 4 AAHC at GFP peak time points) but now using EEG/MEG signals effectively sampled at 200 Hz (i.e., downsampling of 100 Hz by moving-window averaging with 50% overlap).

Figure S1 shows the spatial topographies of EEG and MEG microstates derived at the two sampling rates. We observe a high spatial correspondence among microstate pairs, and in fact the lowest spatial correlation was *R* = 0.83. Therefore, increasing the sampling rate from 40 Hz to 200 Hz has no effect on the spatial signature of microstates.

**Figure S1:**
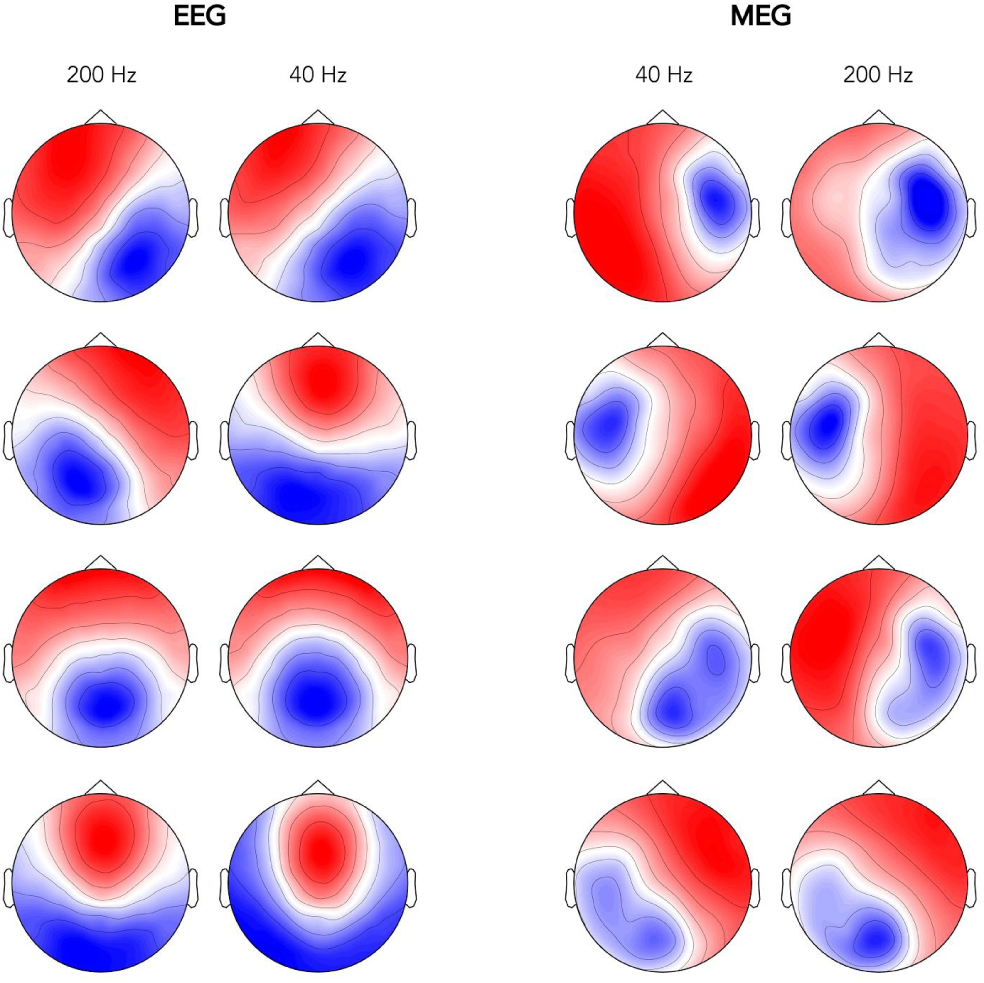
Effect of the sampling rate (40 Hz vs. 200 Hz) on the spatial signature of EEG (**left**) and MEG (**right**) microstates (four-cluster AAHC at GFP peaks). Microstates were paired based on their unambiguous spatial correspondence.

On the other hand, increasing the sampling rate from 40 Hz to 200 Hz significantly decreased the mean lifetime of microstates estimated from their (non-smoothed) activation time series (*t*_41_ = 131.5, *p* = 0 ; compare Table S1 to Table 1 in the main text).

**Table S1:**
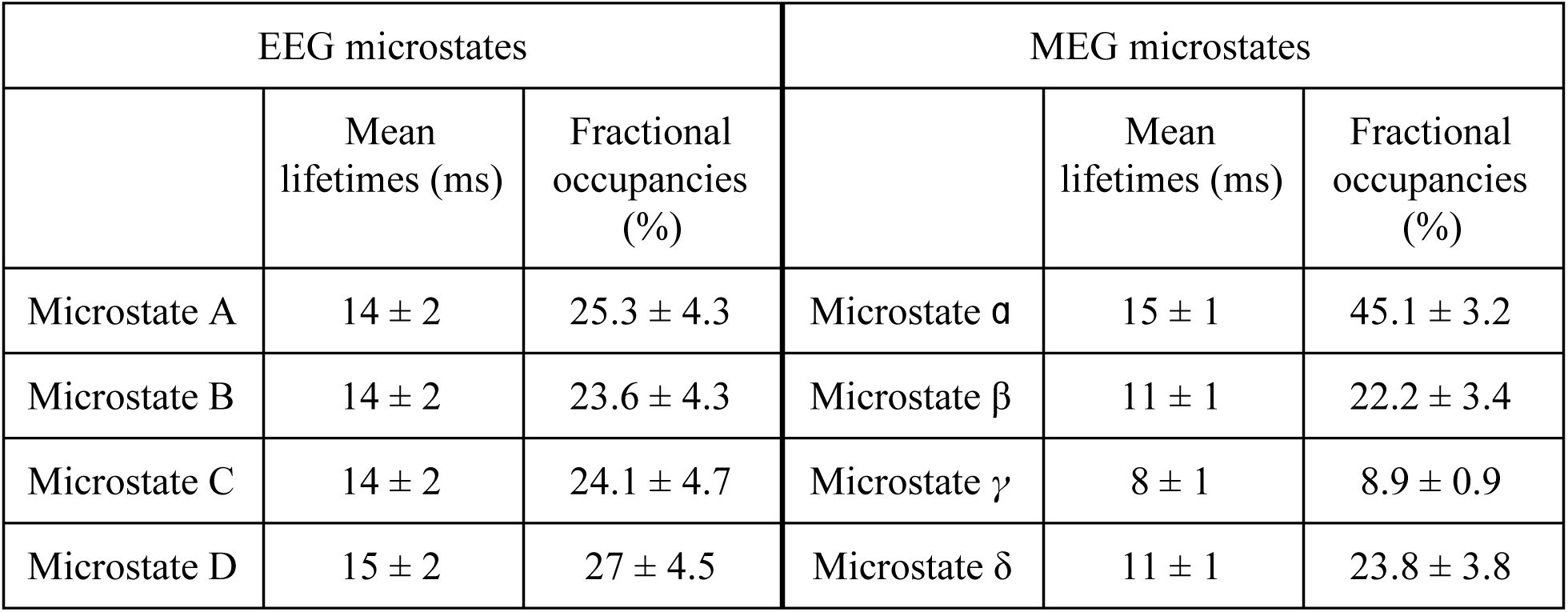
Mean lifetimes and fractional occupancies (mean ± SD) associated with each microstate inferred from EEG or MEG topographies at 200 Hz sampling rate (without temporal smoothing on microstate activation time series).

### S2. Comparison of “two-level” and “group-level” microstate clustering

We repeat here the main microstate analysis (*K* = 4 AAHC of sensor maps downsampled at 40 Hz and restricted to time points of locally maximal GFP), with the only difference that the clustering is directly performed at the group level (i.e., after concatenation of EEG/MEG signals across all subjects), which is more comparable to the group HMM analysis than the two-step microstate clustering presented in the main text.

Figure S2 displays pairs of microstates obtained in both cases and shows that this methodological detail does not impact microstate topographies (spatial correlation among pairs: *R* > 0.71). Mean lifetimes and fractional occupancies were not significantly affected either (*t*_41_ < 1.8, *p* > 0.08 for mean lifetime; *t*_41_ < 1.14, *p* > 0.26 for mean fractional occupancy).

**Figure S2:**
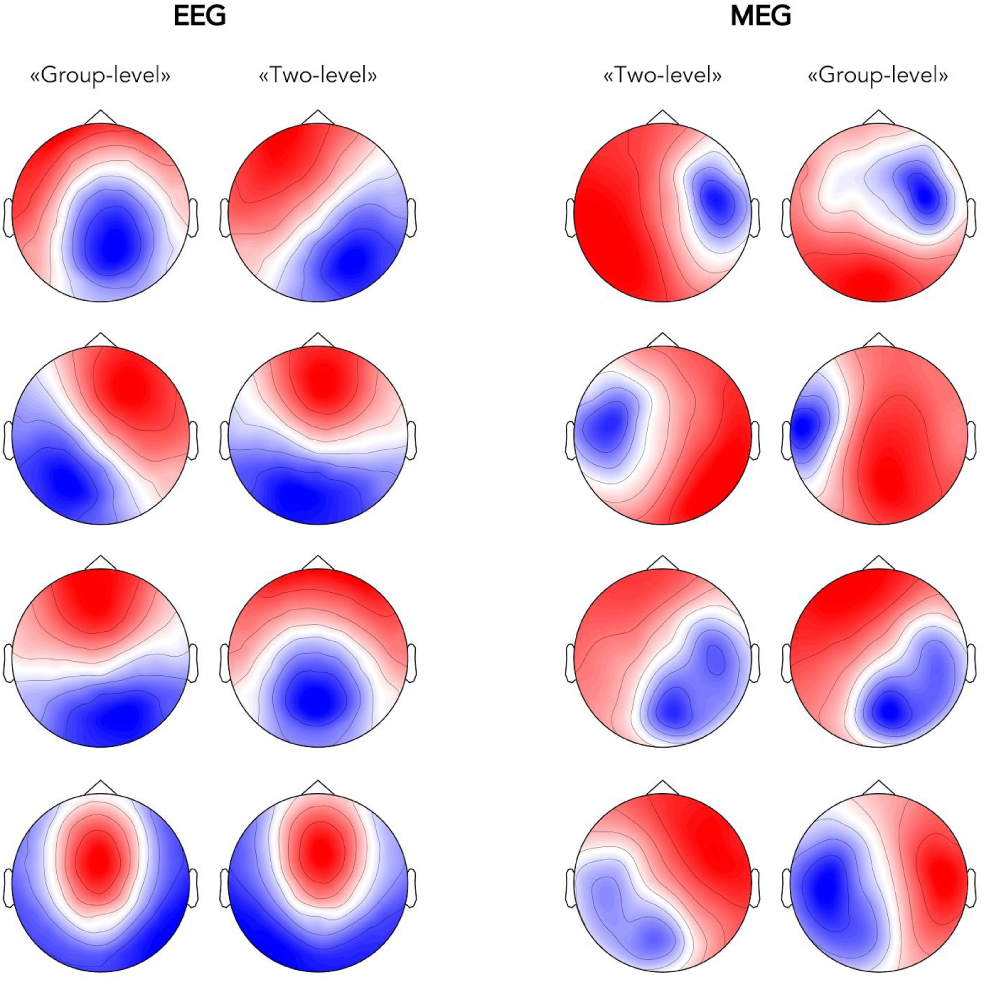
Effect of clustering type (“group-level” vs. “two-level”) on the spatial signature of EEG (**left**) and MEG (**right**) microstates (four-cluster AAHC at GFP peaks, 40 Hz sampling rate). Microstates were paired based on their unambiguous spatial correspondence.

### S3. Microstate clustering without temporal restriction

In the main text, microstate topographies were inferred from local GFP maxima, whereas the HMM ran over the continuous signals. We repeat here the main microstate analysis (*K* = 4 AAHC of sensor maps at 40 Hz sampling rate) but without restriction to isolated time points.

Figure S3 shows that microstate topography is qualitatively unaffected by this methodological difference. As discussed in the main text, this is because the AAHC algorithm explicitly biases microstates towards maximal GFP, suggesting that microstates are mostly sensitive to neural activity around GFP peak times. In line with this suggestion, we observed an excellent one-to-one temporal correspondence between each pair of microstates (Fig. S3).

**Figure S3:**
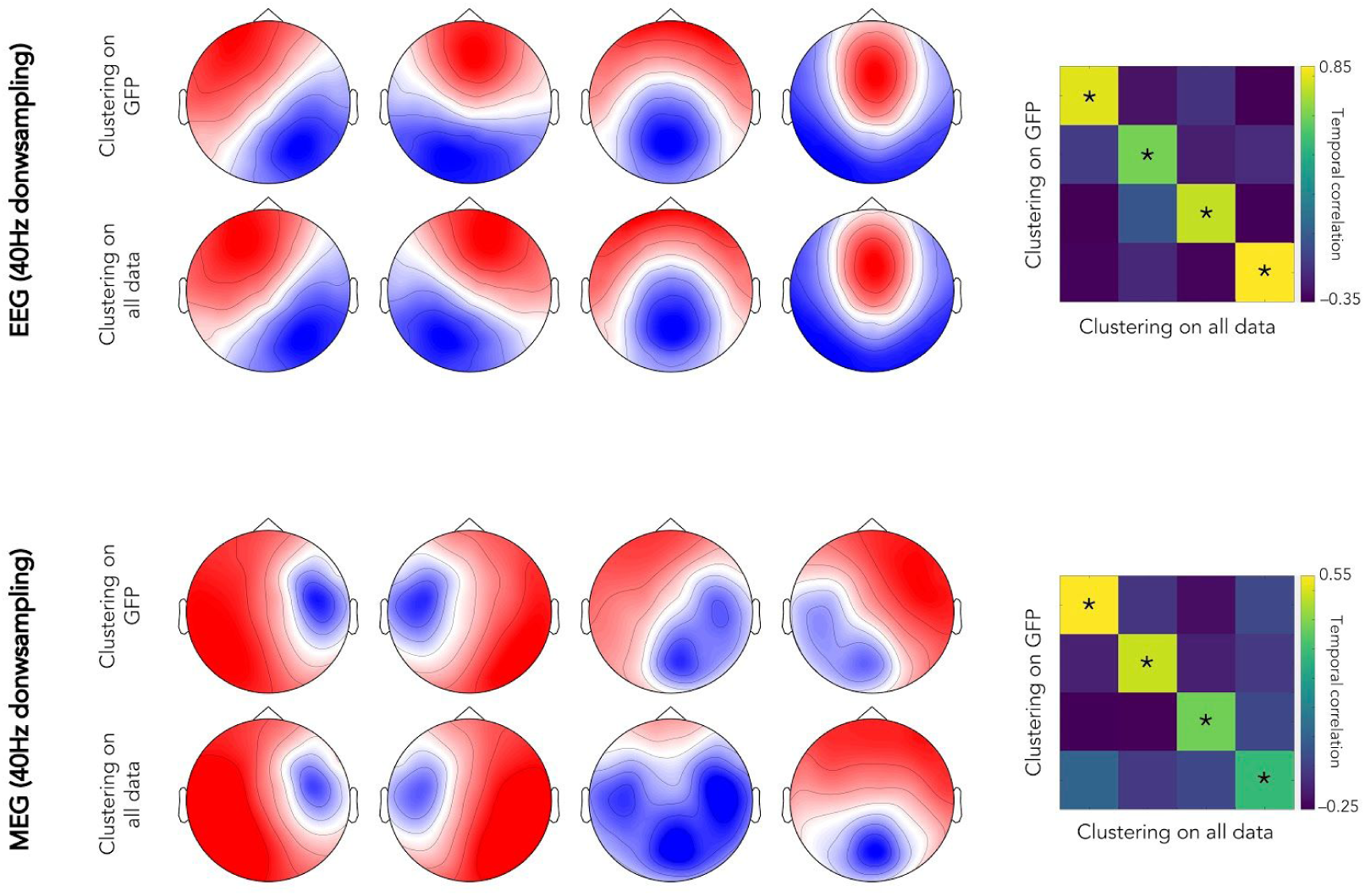
Effect of the temporal restriction to GFP local maxima on EEG (**top**) and MEG (**bottom**) microstates (four-cluster AAHC, 40 Hz sampling rate). Microstates were paired modality based on their unambiguous spatial correspondence. Temporal correlations of microstate activation time series (without temporal smoothing) are shown on the far right.

### S4. Hidden Markov modeling of sensor-versus source-level power envelopes

We focused in the main text on HMM states inferred from sensor power envelopes, but the HMM analysis was developed with, and is commonly applied to, source-level MEG signals. So, we repeated the exact same HMM analysis after source reconstruction. In a nutshell, we applied HMM inference directly on the source power envelope signals used in the main text for the construction of state brain maps. Individual datasets of source envelope signals were demeaned and normalized by the global variance across all sources, and then temporally concatenated across subjects. The resulting dataset was pre-whitened and dimensionally reduced with principal component analysis to retain about 55% of explained variance (leading to *N* = 10 components for EEG and *N* = 53 for MEG). The HMM optimization algorithm was run ten times on these *N*-dimensional time series, each with different initial conditions, and the model with lowest free energy was retained. Brain maps of power increases/decreases upon state activation were computed from the partial correlation between binary state time series (obtained with the Viterbi algorithm) and the group-concatenated source power envelopes.

Figure S4 illustrates the resulting brain maps and shows good spatial agreement between power envelope HMM states inferred from sensor signals and those obtained from reconstructed source activity. Only one state failed to provide a good correspondence with state 1_EEG_/1_MEG_ described in the main text (see top row in Fig. S4). It is also noteworthy that source-level HMM states tended to exhibit more dynamic competition (i.e., including both power increases and decreases).

**Figure S4:**
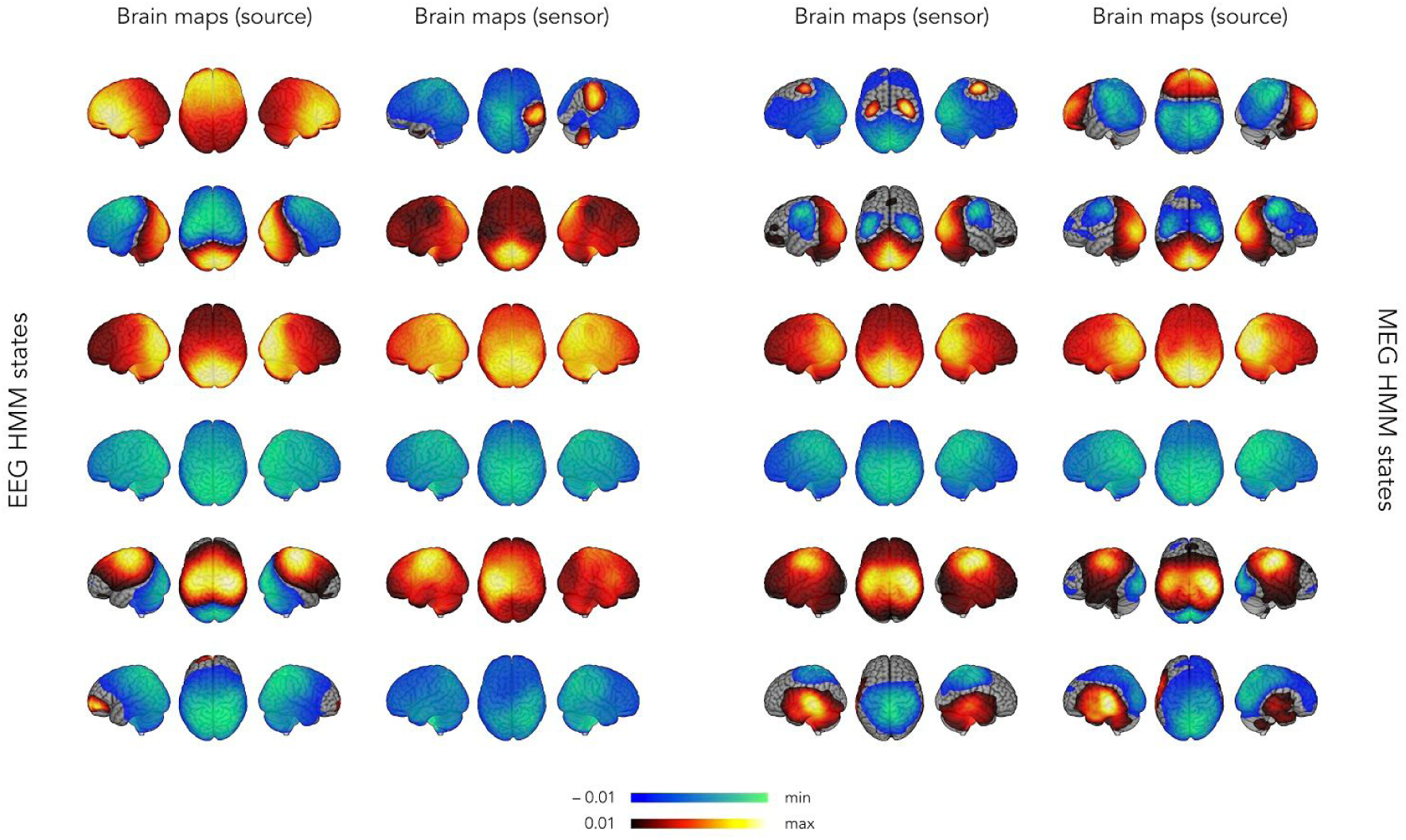
Comparison of HMM states inferred from sensor- and source-level EEG (**left**) and MEG (**right**) power envelopes. Brain maps are thresholded statistically and the lower/upper scales are adapted to the minimum/maximum values. States were ordered based on the visual correspondence.

### S5. Hidden Markov modeling of EEG power envelopes at higher dimensionality

In our main HMM analyses, we fixed the data dimensionality (i.e., the number *N* of principal components fed to the HMM classification algorithm) so the fraction of explained variance is identical for MEG and EEG power envelope signals. This approach naturally takes into account the inherently distinct spatial smoothness of the two recording modalities. However, this difference might impact HMM state classification in EEG and underlie some discrepancies with MEG HMM states. Here, we consider the HMM analysis of EEG power envelopes with *N* = 41, as for MEG.

Figure S5 shows side-by-side the spatial signature of EEG power envelope HMM states at low (*N* = 10; see Fig. 2, left) and higher (*N* = 41; see Fig. 2, right) dimensions. Sensor-level maps show that HMM states are qualitatively similar, although some brain maps exhibited a higher degree of bilaterality (see, e.g., state 1) and more dynamical competition (state 2) when *N* = 41. State 5 was the only state qualitatively different as it involved a frontal activation at when *N* = 41 instead of a left sensorimotor activation at when *N* = 10.

Figure S6 shows the spatial (Fig. S6, top) and temporal (Fig. S6, bottom) correlation analyses between EEG HMM states inferred at *N* = 41 and MEG HMM states at the same dimensionality (Fig. S6, left) or EEG microstates (Fig. S6, right). Despite some qualitative topographical differences, increasing the dimensionality of EEG power envelope data did not affect substantially the spatial or temporal correspondence across state clustering methods or recording modality (compare to Fig. 3). The observations and conclusions discussed in the main text thus stand.

**Figure S5:**
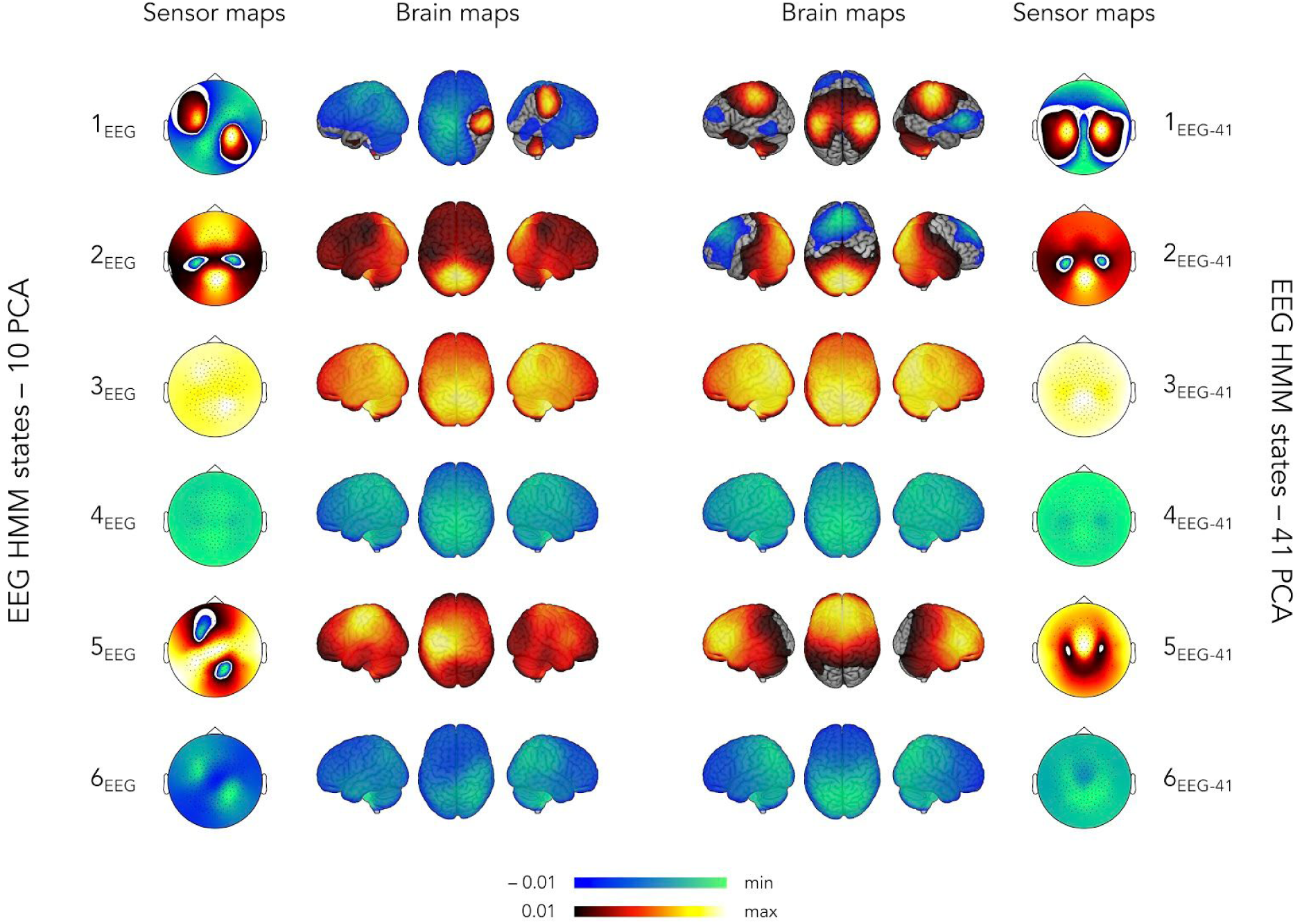
Spatial signature of EEG sensor-level power envelope HMM states obtained with *N* = 10 (**left;** see Fig. 2, left in the main text) and *N* = 41 (**right**) components retained prior to HMM inference.

**Figure S6:**
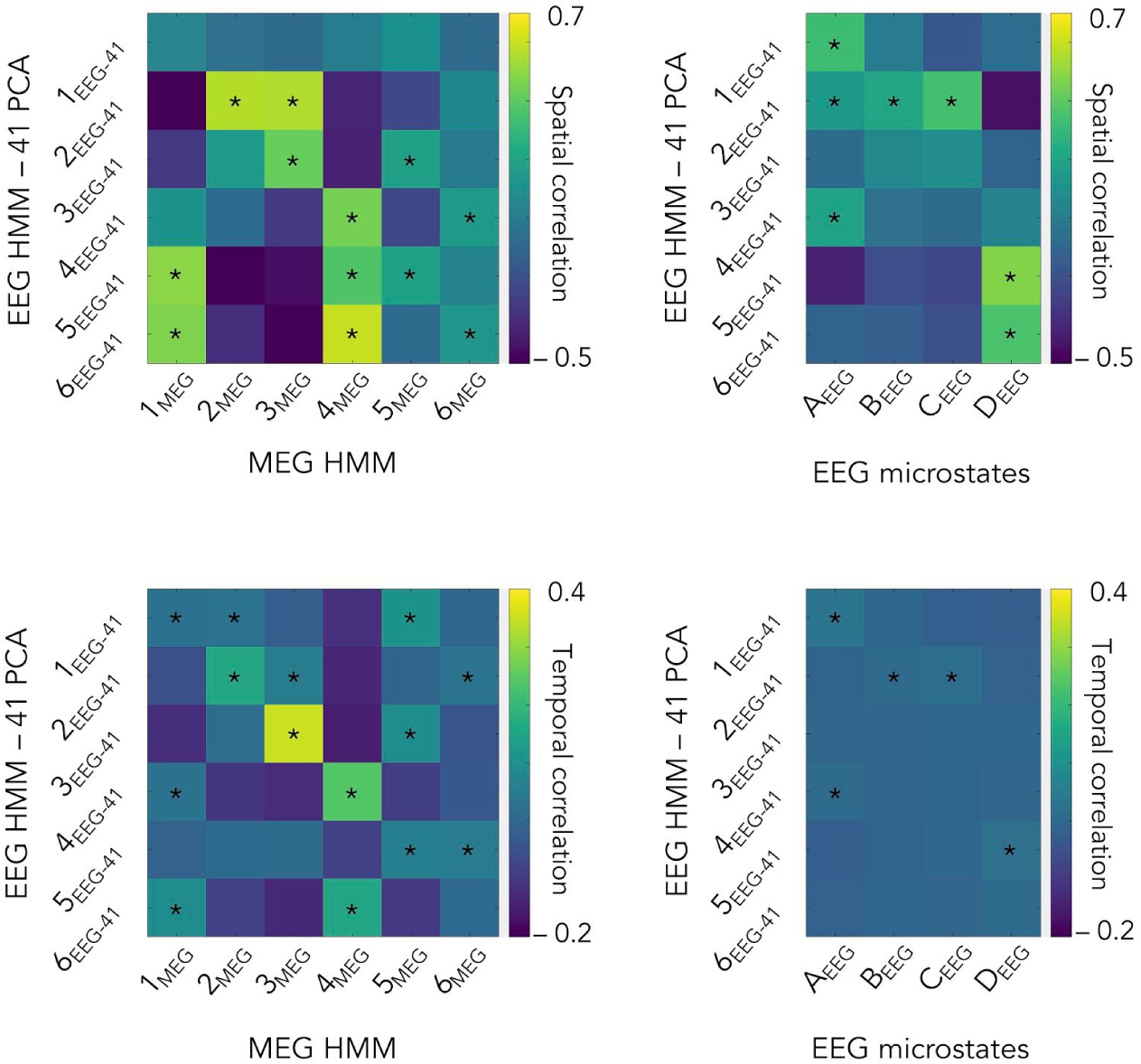
Spatial (**top**) and temporal (**bottom**) correlations when the HMM of EEG power envelopes is inferred from *N* = 41 components. Compare with Fig. 3 in the main text.

### S6. Power envelope hidden Markov model with four states

Finally, we repeated the sensor-level power envelope HMM analysis by lowering the number of states to classify from *K* = 6 (used in the main text) to *K* = 4, for better comparability with the four-microstate clustering.

Figures S7 and S8 present the main results of this analysis. The four-state HMMs merely disclosed a subset of the six-state HMM, specifically states 2_MEG_–5_MEG_ for MEG and to states 1_EEG_, 3_EEG_–5_EEG_ for EEG (compare Fig. S7 to Fig. 2 in the main text). The spatio-temporal correlation analysis comparing microstates vs. HMM states and EEG vs. MEG then naturally led to similar observations than in the case of six-state HMM. The observations and conclusions discussed in the main text thus stand.

**Figure S7:**
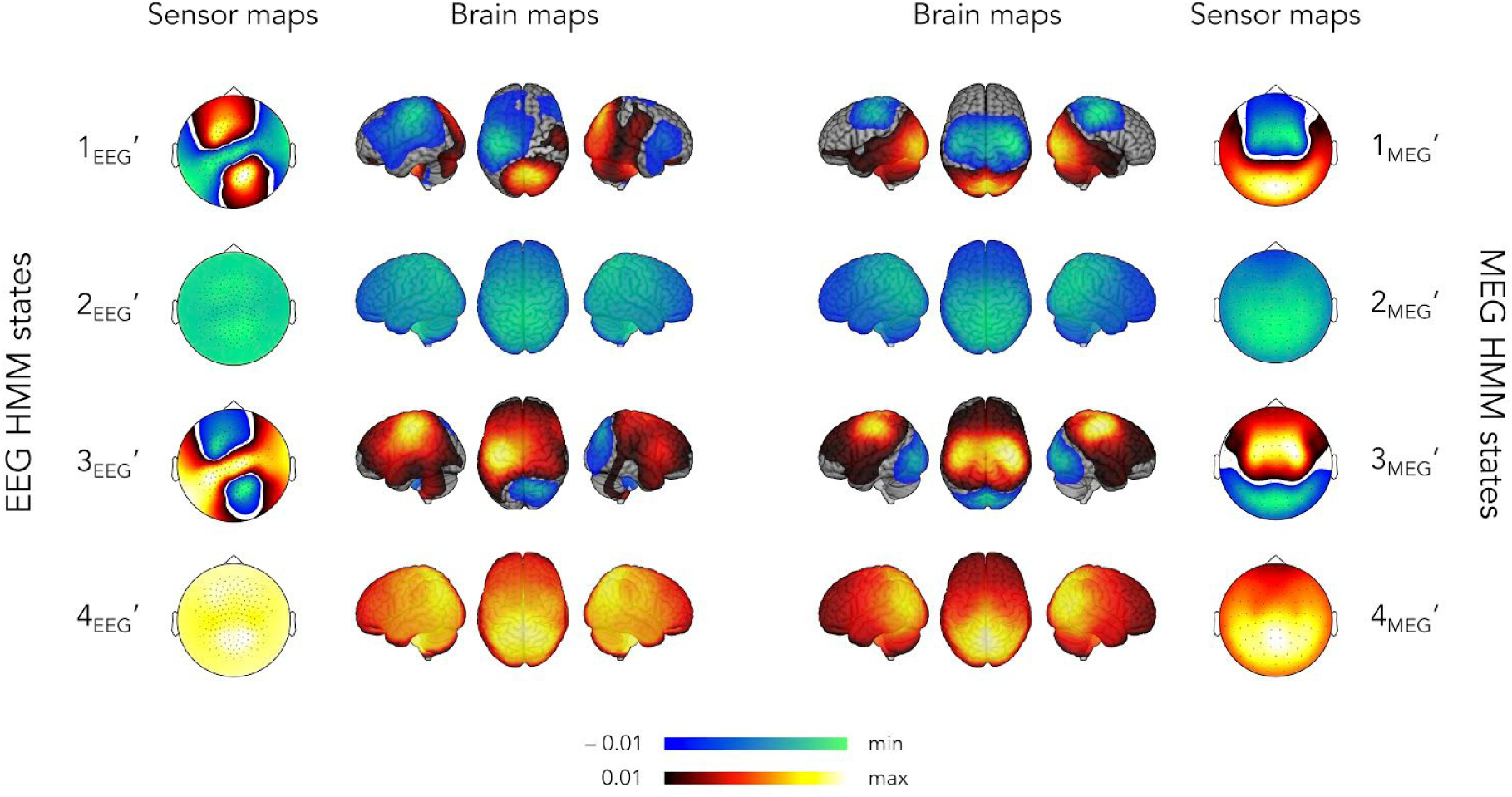
Spatial signature of EEG (**left**) and MEG (**right**) sensor-level power envelope HMM states obtained with *K* = 4. Compare with Fig. 2 in the main text.

**Figure S8:**
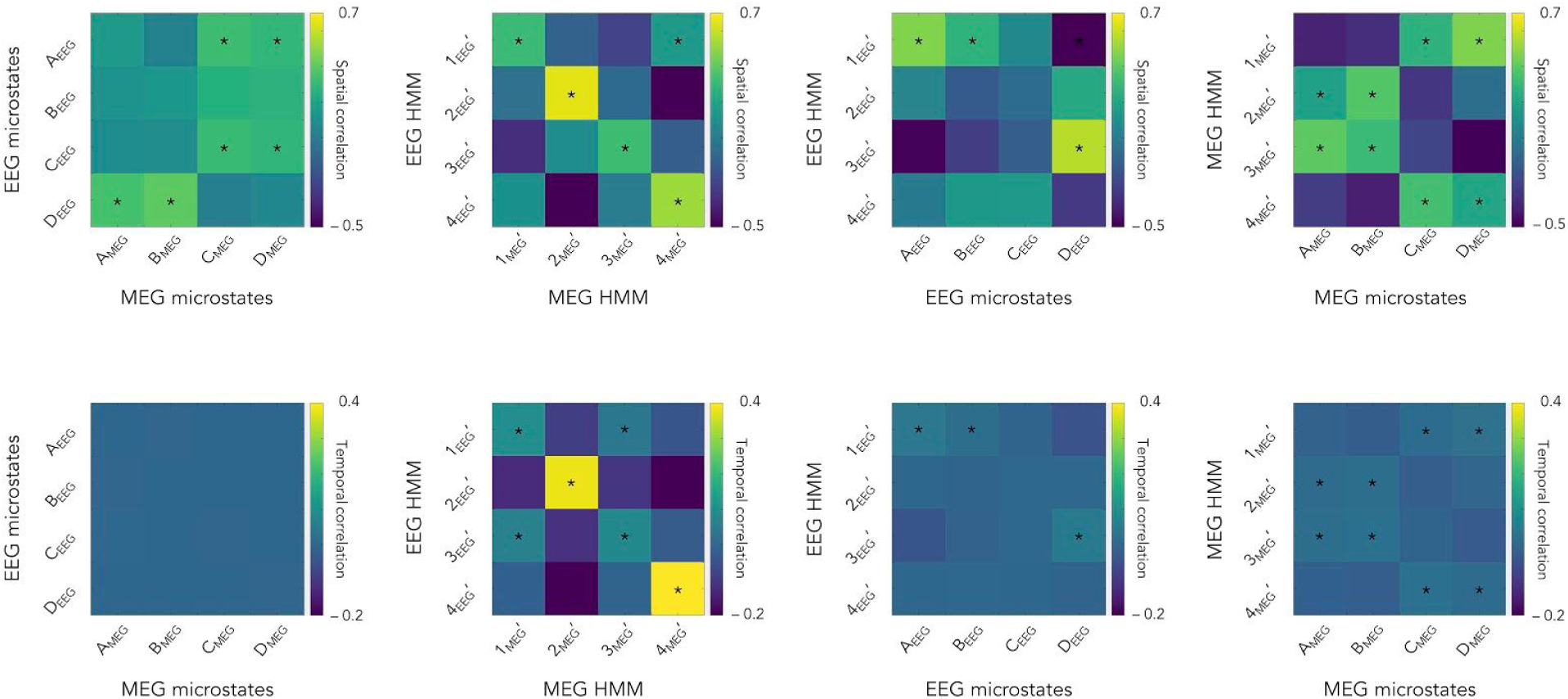
Spatial (**top**) and temporal (**bottom**) correlations in the case of a four-state power envelope HMM. Compare with Fig. 3 in the main text.

## Notes

### Competing Interest Statement

The authors have declared no competing interest.

## References

Baillet, S., 2017. Magnetoencephalography for brain electrophysiology and imaging. Nat. Neurosci. 20, 327–339. https://doi.org/10.1038/nn.4504

Baker, A.P., Brookes, M.J., Rezek, I.A., Smith, S.M., Behrens, T., Probert Smith, P.J., Woolrich, M., 2014. Fast transient networks in spontaneous human brain activity. Elife 3, e01867. https://doi.org/10.7554/eLife.01867

Bertrand, O., Perrin, F., Pernier, J., 1985. A theoretical justification of the average reference in topographic evoked potential studies. Electroencephalogr Clin Neurophysiol 62, 462–464. https://doi.org/10.1016/0168-5597(85)90058-9

Britz, J., Van De Ville, D., Michel, C.M., 2010. BOLD correlates of EEG topography reveal rapid resting-state network dynamics. Neuroimage 52, 1162–1170. https://doi.org/10.1016/j.neuroimage.2010.02.052

Brookes, M.J., Groom, M.J., Liuzzi, L., Hill, R.M., Smith, H.J.F., Briley, P.M., Hall, E.L., Hunt, B.A.E., Gascoyne, L.E., Taylor, M.J., Liddle, P.F., Morris, P.G., Woolrich, M.W., Liddle, E.B., 2018. Altered temporal stability in dynamic neural networks underlies connectivity changes in neurodevelopment. Neuroimage 174, 563–575. https://doi.org/10.1016/j.neuroimage.2018.03.008

Brookes, M.J., Woolrich, M., Luckhoo, H., Price, D., Hale, J.R., Stephenson, M.C., Barnes, G.R., Smith, S.M., Morris, P.G., 2011. Investigating the electrophysiological basis of resting state networks using magnetoencephalography. Proc. Natl. Acad. Sci. U.S.A. 108, 16783–16788. https://doi.org/10.1073/pnas.1112685108

Brunet, D., Murray, M.M., Michel, C.M., 2011. Spatiotemporal Analysis of Multichannel EEG: CARTOOL. Computational Intelligence and Neuroscience. https://doi.org/10.1155/2011/813870

Buzsáki, G., Anastassiou, C.A., Koch, C., 2012. The origin of extracellular fields and currents--EEG, ECoG, LFP and spikes. Nat Rev Neurosci 13, 407–420. https://doi.org/10.1038/nrn3241

Coquelet, N., De Tiège, X., Destoky, F., Roshchupkina, L., Bourguignon, M., Goldman, S., Peigneux, P., Wens, V., 2020a. Comparing MEG and high-density EEG for intrinsic functional connectivity mapping. Neuroimage 210, 116556. https://doi.org/10.1016/j.neuroimage.2020.116556

Coquelet, N., Wens, V., Mary, A., Niesen, M., Puttaert, D., Ranzini, M., Vander Ghinst, M., Bourguignon, M., Peigneux, P., Goldman, S., Woolrich, M., De Tiège, X., 2020b. Changes in electrophysiological static and dynamic human brain functional architecture from childhood to late adulthood. Sci Rep 10, 18986. https://doi.org/10.1038/s41598-020-75858-0

da Cruz, J.R., Favrod, O., Roinishvili, M., Chkonia, E., Brand, A., Mohr, C., Figueiredo, P., Herzog, M.H., 2020. EEG microstates are a candidate endophenotype for schizophrenia. Nat Commun 11, 3089. https://doi.org/10.1038/s41467-020-16914-1

Dale, A.M., Sereno, M.I., 1993. Improved Localization of Cortical Activity by Combining EEG and MEG with MRI Cortical Surface Reconstruction: A Linear Approach. J Cogn Neurosci 5, 162–176. https://doi.org/10.1162/jocn.1993.5.2.162

D’Croz-Baron, D.F., Baker, M., Michel, C.M., Karp, T., 2019. EEG Microstates Analysis in Young Adults With Autism Spectrum Disorder During Resting-State. Front Hum Neurosci 13, 173. https://doi.org/10.3389/fnhum.2019.00173

de Pasquale, F., Della Penna, S., Snyder, A.Z., Marzetti, L., Pizzella, V., Romani, G.L., Corbetta, M., 2012. A cortical core for dynamic integration of functional networks in the resting human brain. Neuron 74, 753–764. https://doi.org/10.1016/j.neuron.2012.03.031

de Pasquale, F., Della Penna, S., Sporns, O., Romani, G.L., Corbetta, M., 2016. A Dynamic Core Network and Global Efficiency in the Resting Human Brain. Cereb. Cortex 26, 4015–4033. https://doi.org/10.1093/cercor/bhv185

De Tiège, X., Op de Beeck, M., Funke, M., Legros, B., Parkkonen, L., Goldman, S., Van Bogaert, P., 2008. Recording epileptic activity with MEG in a light-weight magnetic shield. Epilepsy Res. 82, 227–231. https://doi.org/10.1016/j.eplepsyres.2008.08.011

Della Penna, S., Corbetta, M., Wens, V., de Pasquale, F., 2019. The Impact of the Geometric Correction Scheme on MEG Functional Topology at Rest. Front Neurosci 13, 1114. https://doi.org/10.3389/fnins.2019.01114

Delorme, A., Makeig, S., 2004. EEGLAB: an open source toolbox for analysis of single-trial EEG dynamics including independent component analysis. J Neurosci Methods 134, 9–21. https://doi.org/10.1016/j.jneumeth.2003.10.009

Garcés, P., López-Sanz, D., Maestú, F., Pereda, E., 2017. Choice of Magnetometers and Gradiometers after Signal Space Separation. Sensors (Basel) 17. https://doi.org/10.3390/s17122926

Gschwind, M., Hardmeier, M., Van De Ville, D., Tomescu, M.I., Penner, I.-K., Naegelin, Y., Fuhr, P., Michel, C.M., Seeck, M., 2016. Fluctuations of spontaneous EEG topographies predict disease state in relapsing-remitting multiple sclerosis. Neuroimage Clin 12, 466–477. https://doi.org/10.1016/j.nicl.2016.08.008

Hari, R., Puce, A., 2017. MEG-EEG Primer. Oxford University Press, Oxford, New York.

Hari, R., Salmelin, R., 1997. Human cortical oscillations: a neuromagnetic view through the skull. Trends Neurosci 20, 44–49. https://doi.org/10.1016/S0166-2236(96)10065-5

Higgins, C., Liu, Y., Vidaurre, D., Kurth-Nelson, Z., Dolan, R., Behrens, T., Woolrich, M., 2020. Replay bursts in humans coincide with activation of the default mode and parietal alpha networks. Neuron. https://doi.org/10.1016/j.neuron.2020.12.007

Hipp, J.F., Hawellek, D.J., Corbetta, M., Siegel, M., Engel, A.K., 2012. Large-scale cortical correlation structure of spontaneous oscillatory activity. Nat. Neurosci. 15, 884–890. https://doi.org/10.1038/nn.3101

Hunyadi, B., Woolrich, M.W., Quinn, A.J., Vidaurre, D., De Vos, M., 2019. A dynamic system of brain networks revealed by fast transient EEG fluctuations and their fMRI correlates. NeuroImage 185, 72–82. https://doi.org/10.1016/j.neuroimage.2018.09.082

Kantz, H., Schreiber, T., 2003. Nonlinear Time Series Analysis, 2nd ed. Cambridge University Press, Cambridge. https://doi.org/10.1017/CBO9780511755798

Khanna, A., Pascual-Leone, A., Michel, C.M., Farzan, F., 2015. Microstates in resting-state EEG: current status and future directions. Neurosci Biobehav Rev 49, 105–113. https://doi.org/10.1016/j.neubiorev.2014.12.010

Klimesch, W., 2012. Alpha-band oscillations, attention, and controlled access to stored information. Trends in Cognitive Sciences 16, 606–617. https://doi.org/10.1016/j.tics.2012.10.007

Klimesch, W., Freunberger, R., Sauseng, P., 2010. Oscillatory mechanisms of process binding in memory. Neurosci Biobehav Rev 34, 1002–1014. https://doi.org/10.1016/j.neubiorev.2009.10.004

Koenig, T., Lehmann, D., Merlo, M.C., Kochi, K., Hell, D., Koukkou, M., 1999. A deviant EEG brain microstate in acute, neuroleptic-naive schizophrenics at rest. Eur Arch Psychiatry Clin Neurosci 249, 205–211. https://doi.org/10.1007/s004060050088

Kothe, C.A., Makeig, S., 2013. BCILAB: a platform for brain-computer interface development. J Neural Eng 10, 056014. https://doi.org/10.1088/1741-2560/10/5/056014

Krylova, M., Alizadeh, S., Izyurov, I., Teckentrup, V., Chang, C., van der Meer, J., Erb, M., Kroemer, N., Koenig, T., Walter, M., Jamalabadi, H., 2020. Evidence for modulation of EEG microstate sequence by vigilance level. Neuroimage 224, 117393. https://doi.org/10.1016/j.neuroimage.2020.117393

Lehmann, D., Faber, P.L., Galderisi, S., Herrmann, W.M., Kinoshita, T., Koukkou, M., Mucci, A., Pascual-Marqui, R.D., Saito, N., Wackermann, J., Winterer, G., Koenig, T., 2005. EEG microstate duration and syntax in acute, medication-naive, first-episode schizophrenia: a multi-center study. Psychiatry Res 138, 141–156. https://doi.org/10.1016/j.pscychresns.2004.05.007

Lehmann, D., Ozaki, H., Pal, I., 1987. EEG alpha map series: brain micro-states by space-oriented adaptive segmentation. Electroencephalogr Clin Neurophysiol 67, 271–288.

Lehmann, D., Strik, W.K., Henggeler, B., Koenig, T., Koukkou, M., 1998. Brain electric microstates and momentary conscious mind states as building blocks of spontaneous thinking: I. Visual imagery and abstract thoughts. Int J Psychophysiol 29, 1–11. https://doi.org/10.1016/s0167-8760(97)00098-6

Liu, Q., Farahibozorg, S., Porcaro, C., Wenderoth, N., Mantini, D., 2017. Detecting large-scale networks in the human brain using high-density electroencephalography. Hum Brain Mapp 38, 4631–4643. https://doi.org/10.1002/hbm.23688

Michel, C.M., Brunet, D., 2019. EEG Source Imaging: A Practical Review of the Analysis Steps. Front Neurol 10, 325. https://doi.org/10.3389/fneur.2019.00325

Michel, C.M., Koenig, T., 2018. EEG microstates as a tool for studying the temporal dynamics of whole-brain neuronal networks: A review. Neuroimage 180, 577–593. https://doi.org/10.1016/j.neuroimage.2017.11.062

Michel, C.M., Koenig, T., Brandeis, D., 2009. Electrical neuroimaging in the time domain, in: Michel, C.M., Brandeis, D., Wackermann, J., Gianotti, L.R.R., Koenig, T. (Eds.), Electrical Neuroimaging. Cambridge University Press, Cambridge, pp. 111–144. https://doi.org/10.1017/CBO9780511596889.007

Mishra, A., Englitz, B., Cohen, M.X., 2020. EEG microstates as a continuous phenomenon. Neuroimage 208, 116454. https://doi.org/10.1016/j.neuroimage.2019.116454

Murray, M.M., Brunet, D., Michel, C.M., 2008. Topographic ERP analyses: a step-by-step tutorial review. Brain Topogr 20, 249–264. https://doi.org/10.1007/s10548-008-0054-5

Musso, F., Brinkmeyer, J., Mobascher, A., Warbrick, T., Winterer, G., 2010. Spontaneous brain activity and EEG microstates. A novel EEG/fMRI analysis approach to explore resting-state networks. Neuroimage 52, 1149–1161. https://doi.org/10.1016/j.neuroimage.2010.01.093

Naeije, G., Wens, V., Coquelet, N., Sjøgård, M., Goldman, S., Pandolfo, M., De Tiège, X.P., 2020. Age of onset determines intrinsic functional brain architecture in Friedreich ataxia. Ann Clin Transl Neurol 7, 94–104. https://doi.org/10.1002/acn3.50966

Oldfield, R.C., 1971. The assessment and analysis of handedness: the Edinburgh inventory. Neuropsychologia 9, 97–113.

O’Neill, G.C., Tewarie, P., Vidaurre, D., Liuzzi, L., Woolrich, M.W., Brookes, M.J., 2018. Dynamics of large-scale electrophysiological networks: A technical review. Neuroimage 180, 559–576. https://doi.org/10.1016/j.neuroimage.2017.10.003

Pascual-Marqui, R.D., Michel, C.M., Lehmann, D., 1995. Segmentation of brain electrical activity into microstates: model estimation and validation. IEEE Trans Biomed Eng 42, 658–665. https://doi.org/10.1109/10.391164

Perrin, F., Pernier, J., Bertrand, O., Echallier, J.F., 1989. Spherical splines for scalp potential and current density mapping. Electroencephalogr Clin Neurophysiol 72, 184–187.

Pfurtscheller, G., Lopes da Silva, F.H., 1999. Event-related EEG/MEG synchronization and desynchronization: basic principles. Clin Neurophysiol 110, 1842–1857. https://doi.org/10.1016/s1388-2457(99)00141-8

Puttaert, D., Coquelet, N., Wens, V., Peigneux, P., Fery, P., Rovai, A., Trotta, N., Sadeghi, N., Coolen, T., Bier, J.-C., Goldman, S., De Tiège, X., 2020. Alterations in resting-state network dynamics along the Alzheimer’s disease continuum. In press.

Quinn, A.J., Vidaurre, D., Abeysuriya, R., Becker, R., Nobre, A.C., Woolrich, M.W., 2018. Task-Evoked Dynamic Network Analysis Through Hidden Markov Modeling. Front. Neurosci. 12. https://doi.org/10.3389/fnins.2018.00603

Rabiner, L.R., 1989. A tutorial on hidden Markov models and selected applications in speech recognition. Proc. IEEE 77, 257–286. https://doi.org/10.1109/5.18626.

Rezek, I., Roberts, S., 2005. Ensemble Hidden Markov Models with Extended Observation Densities for Biosignal Analysis, in: Probabilistic Modeling in Bioinformatics and Medical Informatics. Springer-Verlag, pp. 419–450.

Schoffelen, J.-M., Gross, J., 2009. Source connectivity analysis with MEG and EEG. Hum Brain Mapp 30, 1857–1865. https://doi.org/10.1002/hbm.20745

Seedat, Z.A., Quinn, A.J., Vidaurre, D., Liuzzi, L., Gascoyne, L.E., Hunt, B.A.E., O’Neill, G.C., Pakenham, D.O., Mullinger, K.J., Morris, P.G., Woolrich, M.W., Brookes, M.J., 2020. The role of transient spectral “bursts” in functional connectivity: A magnetoencephalography study. Neuroimage 209, 116537. https://doi.org/10.1016/j.neuroimage.2020.116537

Siegel, M., Donner, T.H., Engel, A.K., 2012. Spectral fingerprints of large-scale neuronal interactions. Nat Rev Neurosci 13, 121–134. https://doi.org/10.1038/nrn3137

Siems, M., Pape, A.-A., Hipp, J.F., Siegel, M., 2016. Measuring the cortical correlation structure of spontaneous oscillatory activity with EEG and MEG. Neuroimage 129, 345–355. https://doi.org/10.1016/j.neuroimage.2016.01.055

Sikka, A., Jamalabadi, H., Krylova, M., Alizadeh, S., van der Meer, J.N., Danyeli, L., Deliano, M., Vicheva, P., Hahn, T., Koenig, T., Bathula, D.R., Walter, M., 2020. Investigating the temporal dynamics of electroencephalogram (EEG) microstates using recurrent neural networks. Hum Brain Mapp 41, 2334–2346. https://doi.org/10.1002/hbm.24949

Sitnikova, T., Hughes, J.W., Howard, C.M., Stephens, K.A., Woolrich, M., Salat, D.H., 2020. Spontaneous activity changes in large-scale cortical networks in older adults couple to distinct hemodynamic morphology. bioRxiv. https://doi.org/10.1101/2020.05.05.079749

Sitnikova, T.A., Hughes, J.W., Ahlfors, S.P., Woolrich, M.W., Salat, D.H., 2018. Short timescale abnormalities in the states of spontaneous synchrony in the functional neural networks in Alzheimer’s disease. Neuroimage Clin 20, 128–152. https://doi.org/10.1016/j.nicl.2018.05.028

Sjøgård, M., Bourguignon, M., Costers, L., Dumitrescu, A., Coolen, T., Roshchupkina, L., Destoky, F., Bertels, J., Niesen, M., Vander Ghinst, M., Van Schependom, J., Nagels, G., Urbain, C., Peigneux, P., Goldman, S., Woolrich, M.W., De Tiège, X., Wens, V., 2020a. Intrinsic/extrinsic duality of large-scale neural functional integration in the human brain. bioRxiv. https://doi.org/10.1101/2020.04.21.053579

Sjøgård, M., De Tiège, X., Mary, A., Peigneux, P., Goldman, S., Nagels, G., van Schependom, J., Quinn, A.J., Woolrich, M.W., Wens, V., 2019. Do the posterior midline cortices belong to the electrophysiological default-mode network? Neuroimage 200, 221–230. https://doi.org/10.1016/j.neuroimage.2019.06.052

Sjøgård, M., Wens, V., Van Schependom, J., Costers, L., D’hooghe, M., D’haeseleer, M., Woolrich, M., Goldman, S., Nagels, G., De Tiège, X., 2020b. Brain dysconnectivity relates to disability and cognitive impairment in multiple sclerosis. Hum Brain Mapp. https://doi.org/10.1002/hbm.25247

Skrandies, W., 1990. Global field power and topographic similarity. Brain Topogr 3, 137–141. https://doi.org/10.1007/BF01128870

Tagliazucchi, E., Balenzuela, P., Fraiman, D., Chialvo, D.R., 2012. Criticality in large-scale brain FMRI dynamics unveiled by a novel point process analysis. Front Physiol 3, 15. https://doi.org/10.3389/fphys.2012.00015

Takens, F., 1981. Detecting strange attractors in turbulence, in: Rand, D., Young, L.-S. (Eds.), Dynamical Systems and Turbulence, Warwick 1980, Lecture Notes in Mathematics. Springer, Berlin, Heidelberg, pp. 366–381. https://doi.org/10.1007/BFb0091924

Taulu, S., Simola, J., Kajola, M., 2005. Applications of the Signal Space Separation Method. IEEE Transactions on Signal Processing 53, 3359–3372. https://doi.org/10.1109/TSP.2005.853302

Tibshirani, R., Walther, G., 2005. Cluster Validation by Prediction Strength. Journal of Computational and Graphical Statistics 14, 511–528. https://doi.org/10.1198/106186005X59243

van Ede, F., Quinn, A.J., Woolrich, M.W., Nobre, A.C., 2018. Neural Oscillations: Sustained Rhythms or Transient Burst-Events? Trends in Neurosciences 41, 415–417. https://doi.org/10.1016/j.tins.2018.04.004

Van Schependom, J., Vidaurre, D., Costers, L., Sjøgård, M., D’hooghe, M.B., D’haeseleer, M., Wens, V., De Tiège, X., Goldman, S., Woolrich, M., Nagels, G., 2019. Altered transient brain dynamics in multiple sclerosis: Treatment or pathology? Hum Brain Mapp 40, 4789–4800. https://doi.org/10.1002/hbm.24737

Vidaurre, D., Hunt, L.T., Quinn, A.J., Hunt, B.A.E., Brookes, M.J., Nobre, A.C., Woolrich, M.W., 2018. Spontaneous cortical activity transiently organises into frequency specific phase-coupling networks. Nat Commun 9, 2987. https://doi.org/10.1038/s41467-018-05316-z

Vigário, R., Särelä, J., Jousmäki, V., Hämäläinen, M., Oja, E., 2000. Independent component approach to the analysis of EEG and MEG recordings. IEEE Trans Biomed Eng 47, 589–593. https://doi.org/10.1109/10.841330

von Wegner, F., Bauer, S., Rosenow, F., Triesch, J., Laufs, H., 2021. EEG microstate periodicity explained by rotating phase patterns of resting-state alpha oscillations. Neuroimage 224, 117372. https://doi.org/10.1016/j.neuroimage.2020.117372

Wens, V., 2015. Investigating complex networks with inverse models: analytical aspects of spatial leakage and connectivity estimation. Phys Rev E Stat Nonlin Soft Matter Phys 91, 012823. https://doi.org/10.1103/PhysRevE.91.012823

Wens, V., Bourguignon, M., Vander Ghinst, M., Mary, A., Marty, B., Coquelet, N., Naeije, G., Peigneux, P., Goldman, S., De Tiège, X., 2019. Synchrony, metastability, dynamic integration, and competition in the spontaneous functional connectivity of the human brain. Neuroimage 199, 313–324. https://doi.org/10.1016/j.neuroimage.2019.05.081

Wens, V., Marty, B., Mary, A., Bourguignon, M., Op de Beeck, M., Goldman, S., Van Bogaert, P., Peigneux, P., De Tiège, X., 2015. A geometric correction scheme for spatial leakage effects in MEG/EEG seed-based functional connectivity mapping. Hum Brain Mapp 36, 4604–4621. https://doi.org/10.1002/hbm.22943

Wens, V., Mary, A., Bourguignon, M., Goldman, S., Marty, B., Op de Beeck, M., Bogaert, P.V., Peigneux, P., De Tiège, X., 2014. About the electrophysiological basis of resting state networks. Clin Neurophysiol 125, 1711–1713. https://doi.org/10.1016/j.clinph.2013.11.039

Woolrich, M.W., Baker, A., Luckhoo, H., Mohseni, H., Barnes, G., Brookes, M., Rezek, I., 2013. Dynamic state allocation for MEG source reconstruction. Neuroimage 77, 77–92. https://doi.org/10.1016/j.neuroimage.2013.03.036

Yao, D., 2017. Is the Surface Potential Integral of a Dipole in a Volume Conductor Always Zero? A Cloud Over the Average Reference of EEG and ERP. Brain Topogr 30, 161–171. https://doi.org/10.1007/s10548-016-0543-x

Yuan, H., Zotev, V., Phillips, R., Drevets, W.C., Bodurka, J., 2012. Spatiotemporal dynamics of the brain at rest — Exploring EEG microstates as electrophysiological signatures of BOLD resting state networks. NeuroImage 60, 2062–2072. https://doi.org/10.1016/j.neuroimage.2012.02.031

